# The development of social attention in orangutans: comparing peering behaviour in wild and zoo-housed individuals

**DOI:** 10.1101/2024.05.31.596770

**Authors:** Paulina Kukofka, Richard Young, Julia A. Kunz, Lara Nellissen, Shauhin E. Alavi, Fitriah Basalamah, Daniel B. M. Haun, Caroline Schuppli

## Abstract

Social learning is the cornerstone of all cultural processes and plays a pivotal role during the evolution of cognition. To understand how social learning evolved, we have to look at the immediate and developmental conditions affecting individuals’ tendencies to attend to social information. We compared peering behaviour (i.e., close-range and sustained observation of the activities of conspecifics) in wild and zoo-housed immature Sumatran orangutans (*Pongo abelii*) by analysing long-term data which included 3101 peering events collected at the Suaq Balimbing research site in Indonesia and at four European zoos on 35 immature individuals. Using Generalized-Additive-Mixed-Models, we tested for age-specific patterns in peering frequency, target, and context selection. We found similar age trajectories of peering in both settings but higher mean frequencies of peering in the zoos, even after controlling for varying social opportunities to peer. Wild immatures preferably peered at their mothers but zoo-housed immatures at non-mother individuals. In both settings, immatures preferred to peer at older individuals, and in learning-intense contexts. Our findings suggest a hard-wired component in the tendency to attend to social information and a considerable degree of ontogenetic plasticity - a combination that was likely foundational for the evolution of complex cultures, including human culture.

**Highlights:**

- We compared attendance to social information in wild and zoo-housed orangutans
- Immature orangutans peer in contexts where learning is expected
- Peering frequency develops similarly over age, suggesting hardwired propensities
- Peering target and context selection differs between the two settings
- Orangutans’ tendency to attend to social information shows plasticity

## 1. Background

Like humans, many animals are born with an incomplete skill set and thus need to acquire a substantial share of the knowledge and skills required for survival and reproduction during development (Schuppli et al. 2012; Ross and Jones 1999). For this purpose, many species rely on social learning which is “learning that is influenced by observation of, or interaction with, another animal (typically a conspecific) or its products” (Heyes 1994; Heyes 2012; reviewed by Hoppitt and Laland 2013; Whiten and van de Waal 2018). Compared to independent learning, social learning allows for more efficient learning, as individuals can use social cues to guide their exploration, tap into skill sets already present in their population, and consequently avoid the costs and potential risks of innovation (van Schaik and Burkart 2011; Whiten and van Schaik 2007; Watson et al. 2018). In its simplest form, social learning is non-observational, and knowledge is transferred through social facilitation or local enhancement (Whiten et al. 2004; Rapaport and Brown 2008). However, to acquire more learning-intense skills such as tool use, high-fidelity forms of social learning, namely observational social learning (e.g., imitation or emulation) or interactions with other individuals (e.g., teaching) are needed to allow for reliable transmission of information (Carvajal and Schuppli 2022).

The ability to learn socially is likely an adaptive behaviour (van Schaik and Burkart 2011), and social learning is a prerequisite for the formation of traditions and cultures (van Schaik et al. 2003; Whiten et al. 1999; Heyes 2012) which are most pronounced and advanced in great apes, most of all humans (Jaeggi et al. 2010; van Schaik et al. 2017; Whiten et al. 1999; van Schaik et al. 2003). Cultural processes based on high-fidelity forms of social learning (i.e., observational or interactive) have played a pivotal role in the evolution of human cognition (Boyd et al. 2011; Herrmann et al. 2007). Consequently, it is important to understand how social learning, especially high-fidelity social learning, has evolved. So far, most studies infer social learning in wild animals by looking at the results of social learning, such as the distribution of cultural elements across populations or the spread of behaviours through populations (e.g., Whiten et al. 1999; van Schaik et al. 2003). However, social learning happens at the individual, not the population level. As such, attending to information provided by other individuals (i.e., social information) is a crucial element of social learning. So far, it remains unclear what ecological and social conditions affect individuals’ tendencies to attend to social information. Ecological and social conditions can affect attendance to social information on the immediate mechanistic level (i.e., prevailing conditions that increase or decrease attention to social information, henceforward called “immediate level”) and the developmental ontogenetic level (i.e., factors and conditions experienced during development that affect an individual’s current tendency to attend to social information, henceforward called “developmental level”) (Tinbergen 1963). However, to be subject to classic evolutionary processes, the tendency to attend to social information must, at least to some extent, be genetically coded. By looking at the process of individuals attending to social information (Schuppli and van Schaik 2019b), we can identify the immediate and developmental ecological and social conditions that favour social learning and infer heritable predispositions (Afseth et al. 2022). This may lead to novel insights into the evolution of social learning.

Field studies on wild animals allow insights into their behaviour under natural conditions. Comparative studies of individuals and populations that live under different social or ecological conditions, e.g., between wild and captive populations, can help to disentangle extrinsic and intrinsic factors affecting a behaviour (Harrison and van de Waal 2022). Fluctuations in the natural environment, such as food availability, predation risk, or the availability of social partners, make the identification of certain behavioural trends and patterns possible. Yet, pinpointing the exact underlying mechanisms at play often remains difficult. In contrast to wild animals, captive animals live in conditions that are usually far removed from their naturally occurring habitats, and their behaviours can differ greatly from those observed in the wild (Forss et al. 2015; Kummer and Goodall 1985; Benson-Amram et al. 2013). These behavioural differences have been attributed to factors such as excess of energy and free time (Kummer and Goodall 1985; Haslam 2013) or habituation to humans (Forss et al. 2022; Damerius et al. 2017). Captive conditions may help to elucidate the underlying mechanisms of observed behaviour because they can confirm or challenge patterns observed under natural conditions (Palagi and Bergman 2021). Furthermore, by comparing a behaviour observed in the wild to its expression in captivity, we can investigate behavioural flexibility, and disentangle genetic and developmental effects.

Orangutans are well suited for the study of social learning mechanisms because of their high tendency to attend to social information which plays an important role during skill acquisition (van Schaik 1999; Schuppli and van Schaik 2019b; Schuppli et al. 2016b; Jaeggi et al. 2010; Mikeliban et al. 2021; Mörchen et al. 2023). Among the orangutans’ socially learned skills are foraging skills which develop gradually during the dependency period and beyond (Mendonça et al. 2017; Schuppli et al. 2016a). Furthermore, the orangutans’ exceptionally slow development implies an extended period of immaturity and thus drawn-out time for learning. Before weaning they go through a six-to nine-year dependency period during which they stay in constant close proximity to their mother who is their main interaction partner (Schuppli et al. 2016b; Mikeliban et al. 2021; Scott et al. 2023). Orangutans are semi-solitary and have fission-fusion social dynamics (Mitra Setia et al. 2009; van Schaik 1999). Depending on species and population, independent immatures and adults spend between 40 – 90% of their time on their own, meaning they have limited opportunities to attend to social information (van Noordwijk et al. 2009). During the dependency period, learning opportunities from other individuals are limited to the mother’s association partners. The slow development and low level of social interactions during dependency facilitate the tracking and documenting of learning opportunities, which is more challenging in species with a faster developmental period and more associations.

Observational learning through peering, i.e., close-range and sustained observation of a conspecific’s activities (Schuppli et al. 2016b; van Schaik et al. 2016), is the most prominent and best-studied form of social learning that wild orangutans use. Peering has also been described in other primate species, including chimpanzees (Matsuzawa et al. 2001; Lonsdorf 2006), bonobos (Idani 1995; Johnson et al. 1999; Stevens et al. 2005; Stevens et al. 2006), brown capuchin monkeys (Ottoni et al. 2005), white-faced capuchin monkeys (Perry and Jiminez 2012), and vervet monkeys (Grampp et al. 2019). Not all peering may lead to social learning and peering may have additional social functions (Vervaecke et al. 2000; Stevens et al. 2006). However, during peering, individuals attend to social information via sustained, close-range observation of the activities of a conspecific, which is a likely prerequisite for observational social learning. There is strong evidence that overall, peering is indeed a means for social learning: Peering mainly occurs in contexts where learning is expected, such as foraging, nest-building, or social interactions (Schuppli et al. 2016b; van Schaik et al. 2017), and it increases with increasing processing intensity and rarity of a displayed behaviour (Schuppli et al. 2016b; Perry and Jiminez 2012; Ottoni et al. 2005). Furthermore, peering rates are highest during immaturity, when individuals have to learn most of their skills (Schuppli et al. 2016b; Schuppli and van Schaik 2019a; Lonsdorf 2006; Matsuzawa et al. 2001) and peering is often followed by selective practice of the observed skill (Schuppli et al. 2016b; Schuppli and van Schaik 2019a). To date, it remains unclear what factors modulate peering behaviour. The tendency to peer is likely regulated by a mechanism different from conscious desire to learn. Investigating the factors that prompt an individual to engage in peering will increase our understanding of the underlying mechanisms and may shed novel light on the evolution of peering and observational social learning.

Wild orangutans peer up to 40’000 times during their lifetime (Schuppli and van Schaik 2019b) with most peering occurring during immaturity (Schuppli et al. 2017). Orangutans show biases in their peering target selection (i.e., the individuals they peer at) (Schuppli et al. 2016b; Schuppli and van Schaik 2019a). With increasing age, peering preferences shift from the mother to non-mother individuals (Schuppli et al. 2016b), with most non-mother peering first being directed at adults and later at peers (Schuppli and van Schaik 2019a) and with females and males having different peering target preferences (Ehmann et al. 2021). Comparative studies have found that the more gregarious and socially tolerant Sumatran orangutans (*Pongo abelii*) (van Schaik 1999) show higher peering rates than the less gregarious Bornean orangutans (*P. pygmaeus*) (van Schaik et al. 2016; Schuppli et al. 2017), even after controlling for differences in association time (and hence opportunities to peer) and prevailing food availability (Mörchen et al. 2024). This difference suggests that either growing up in more sociable and ecologically favourable conditions increases individuals’ tendency to peer or that the two orangutan species differ in their intrinsic tendency to peer (Schuppli et al. 2017; Schuppli et al. 2020). However, despite these findings, many aspects of how social and ecological conditions shape peering behaviour on the immediate and developmental levels remain unclear. Furthermore, because these studies compared two different populations and species, it remains unclear whether genetic differences cause the observed group-level variation.

In this study, we compared the peering behaviour of wild Sumatran orangutans to peering behaviour of zoo-housed conspecifics to investigate its underlying mechanisms. To do so, we used a detailed comparative longitudinal dataset. We looked at the frequency of peering behaviour across ages, the development of peering targets across ages, and the behavioural contexts in which peering occurred. Zoo-housed and wild orangutans differ in their ecological and social environment. Zoo orangutans are housed in stable social groups, which – unlike for other great apes – stands in stark contrast to their natural social environment and makes them an especially suited species for this study. Social factors such as the number of peering targets available are quantifiable. They can thus be controlled for (Boesch 2021), which we use to pin down whether social or ecological factors are at work. We predicted that:

### 1. Peering as a means for social learning

If zoo-housed orangutans peer to learn, they will selectively peer in learning intense contexts, such as when rare foods are eaten and when pre-ingestive processing is more intense, as has been found for wild orangutans.

### 2. Ontogeny of peering

a. If ecological necessity to learn skills brings about peering behaviour on the immediate or developmental level, zoo-housed orangutans will peer less than their wild conspecifics, as they are provisioned and thus have easy-to-process food abundantly available which reduces the time they spend foraging and the need to learn processing-intense foraging techniques.
b. If peering behaviour is shaped by social opportunities (i.e., the availability of peering targets) on the immediate or developmental level, zoo-housed orangutans will peer more than wild orangutans, because they are in constant and close association with conspecifics.

### 3. Development of peering target selection

a. If peering target selection is shaped by ecological necessity/pressure on the immediate or developmental level to learn from knowledgeable individuals, compared to wild orangutans, zoo-housed orangutans will show no specific patterns in their peering target selection, as all individuals exhibit little variation in knowledge and skill sets.
b. If peering target selection is shaped by social opportunities (i.e., the availability or lack of different peering targets) on the immediate or developmental level, compared to wild orangutans, zoo-housed orangutans will exhibit less bias towards a specific age class, as they have all classes of peering targets readily available.

### 4. Peering context selection

a. If peering context selection is shaped on the immediate or developmental level by the ecological necessity to learn, compared to wild orangutans, zoo-housed orangutans will peer less frequently in the feeding and nest-building context, as these behaviours tend to be less complex in zoos (see Fig. A8). Further, zoo-housed orangutans will show higher peering rates in social contexts since social interactions are more frequent and more complex than in the wild.
b. If peering context selection is shaped on the immediate or developmental level by social opportunities to peer, compared to wild orangutans, zoo-housed orangutans will show less pronounced preferences in their context selection and also peer in contexts that offer less learning input.

## 2. Methods

### 2.1. Data collection

The data from the wild population of Sumatran orangutans (*Pongo abelii*) were collected over 12 years (October 2008 to January 2020) at the Suaq Balimbing monitoring station, located in the Gunung Leuser National Park in South Aceh, Indonesia (3°42’N, 97°26’E). The data on zoo-housed Sumatran orangutans were collected in zoos in Switzerland (Zoo Zürich and Zoo Basel) and Germany (Zoo Leipzig and Zoo Dresden) over the course of two years (May 2021 to July 2023). All data collected for this study were obtained through behavioural observations, following an established protocol for orangutan behavioural data collection (see https://www.ab.mpg.de/571325/standarddatacollectionrules_suaq_detailed_jan204.pdf). This study used two types of data: I) instantaneous scan sampling at two-minute intervals of the individual’s activity, its visibility, and its distance to other orangutans within a 50-meter radius (association partners), and II) all-occurrence sampling of peering behaviour. Both data types were collected simultaneously during focal follows.

Peering is defined as “directly looking at the actions of another individual sustained over at least 5 seconds, and at a close enough range that enables the peering individual to observe the details of the action (within 2 meters in the feeding and within 5 meters in the nest-building context)” (see Schuppli et al. 2016b). At every peering event, details such as the peering target, its activity, and the duration of the peering event were recorded. All observers whose data on wild orangutans were included in this study passed an interobserver reliability test of more than 85 % accordance in simultaneous observations of the same individual. All-occurrence peering data used from Suaq were collected by sixteen observers. All observers in the zoo underwent training before collecting data and were closely supervised during follows to ensure accurate data entry. Overall, we included 8750 hours and 1190 hours of behavioural data on wild and zoo-housed orangutans, respectively, which resulted in 1915 observed peering events in the wild and 1186 peering events in the zoos.

### 2.2. Focal animals

We included observations on 65 orangutans (40 from the wild, 25 from the zoos, Table A1). Thirty-five of the individuals were immatures at the time of data collection, all from known mothers, 22 from the wild (10 females, 12 males) and 13 housed in zoos (9 females, 4 males). The immatures’ age ranged from 0.5 to 15.9 years (mean = 5.8 years) in the wild and from 0.6 to 14.0 years (mean = 5.5 years) in the zoos. As the exact dates of birth for most of the individuals in the wild are unknown, we worked with estimates calculated by experienced observers upon first encounter of the animals. For wild adult females, ages are estimated based on their known number of offspring. The number of offspring is inferred based on observed dependent offspring and genetically confirmed offspring. For adult males with unknown birth dates, we assigned standardized birth dates: for unflanged males twenty years, and for flanged males thirty years before the date on which they were first encountered in the study area (van Noordwijk et al. 2023). For orangutans born in the zoos, exact birth dates are known. Orangutans in zoo environments may differ in the time they reach developmental milestones such as age at first reproduction, but reliable data investigating developmental differences between zoo-housed and wild populations are missing to date. To simplify later analysis, we defined the age limits of each age class based on insights obtained from wild Sumatran orangutans: throughout this study “immatures” refers to all individuals below the age of sixteen years. Individuals below the age of 8 years were classified as “dependent immatures” (based on the fact that at Suaq the average weaning age is 8.1 years, Schuppli unpublished data), and immatures between the age of 8.1 and 16 years as “independent immatures” (van Noordwijk et al. 2009; Wich et al. 2009). None of the female immatures in our data set had reproduced yet but one male at the zoo who was 14 at the time of data collection had sired an offspring.

### 2.3. Data sets and statistical analyses

The full data set included data from 1061 follow days of which 567 were from immature orangutans: 466 days from the wild data and 101 days from the zoo data (17 from Basel, 12 from Dresden, 25 from Leipzig, and 47 from Zurich). *The data sets used for this study will be made available through a publicly accessible data repository upon publication*. The average observation duration on a follow day from immatures in the wild was 9.7 hours (ranging from 0.3 to 13.6 h) and 8.2 hours (ranging from 3.6 to 10.4 h) in the zoos, with 76 % of the wild data, and 89 % of the zoo data being full day follows (i.e., nest-to-nest in the wild, from morning until the animals settle in to sleep at the zoos). We used the all-occurrence data on peering behaviour to analyse peering frequencies, peering target selection, and peering context selection, and the two-minute scan data to calculate overall visible observation duration, as well as time spent in peering range (0 to 2 m) to association partners. Because visibility of a focal individual varied during a follow, we corrected for these differences by excluding scans from our analysis where the focal was out of sight of the observer. In the wild, visibility has only been collected since 2014 but was positively correlated with age of the individual (LME, Estimate (Age 0-10) = 0.038, *p* < 0.001, Figure A1a, Table A2). For data collected before 2014 we therefore used predicted visible observation duration values obtained via the model estimates of a Linear Mixed-Effects Model (Table A2).

All plots and all statistical analyses were performed in R (R Core Team 2023). All models included age (which we z-transformed) as predictor to assess the effects of age on peering behaviour. Visualization of the raw data showed that age had a non-linear effect on most of our response variables. To account for this non-linearity and ensure the best fit for our data, we ran Generalized Additive Mixed Models (GAMM) using the gam function from the mgcv package (Wood 2018). As an extension of a generalized linear model (GLM), GAMM uses smoothing splines to estimate linear and non-linear relationships (Wood 2017). We formulated all models with counts as the response variable with a negative-binomial error distribution to avoid zero inflation issues caused by the large number of zeros in our data sets (as a result of follow days where no peering occurred, thus causing “true” zeros). For models with proportions as the response variable, we used a beta distribution. For all models, we reported the adjusted r2 and deviance explained (DE) as obtained via the model output (see Table A2 for an overview of all model structures).

Since we tested for differences in peering between the wild and the zoo, setting was included as a predictor in all the models (with the exemption of models 1, and A1), as well as an interaction between age and setting, to test if age patterns in peering develop differently between settings. We modelled a smooth of age, and a tensor interaction between age and setting, to account for the non-linearity of these terms. We included individual ID as a random smooth in all models to avoid pseudo replication issues as individuals appeared multiple times in the data sets. To account for potential variation between different observers, we also included observer ID as a random smooth. Statistical significance was assessed at the 5 % level for all analyses. We used the difference_smooths function of the gratia package (Simpson 2024) to visually assess statistical significant differences between the age splines of each setting.

Aside from our main analyses, we tested whether individuals of different zoos significantly differed in their peering rates. Therefore, we fitted a GAM model including age, and site (i.e., Suaq, Zoo Basel, Zoo Dresden, Zoo Leipzig, Zoo Zürich), as fixed factors and individual ID and observer ID as random smooths (Fig. A2, Table A3). Visualization of the data suggests that individuals in the zoos with smaller group sizes (Zoo Basel and Zoo Dresden) show lower peering rates compared to individuals in the zoos with larger group sizes (Zoo Zürich and Zoo Leipzig, Fig. A2). However, post-hoc analysis using the emmeans function from the emmeans package (Lenth 2023) showed that there were no statistically significant differences in peering rates between the different zoos (Table A4). We consequently didn’t distinguish between the zoos in our main analyses.

#### 2.3.1. Peering as a means for social learning

To assess whether peering is used as a means to social learning in dependent immature zoo-housed orangutans, we tested the effects of a food item’s processing intensity and frequency on peering behaviour. We classified food items into difficult and easy to process food items. Difficult food items require multiple steps of processing before ingestion, such as dexterous manipulations to extract edible parts (e.g., peeling, breaking off, insertion of a tool). Easy food items can be ingested directly without prior processing (i.e., pick and eat) (see Jaeggi et al. 2008; Schuppli et al. 2016b). We counted peering events at easy and difficult food items per individual per day. Using the two-minute scan data we calculated the time adult association partners spent feeding on easy and difficult food items on that specific day. We then calculated peering rates controlled for opportunities to peer for each item processing class by dividing the number of peering events by the time adult association partners had spent feeding on it. To obtain a measure of the overall frequency of a food item, we divided the time adult association partners spent feeding on each item by the total visible observation duration. To calculate peering rates controlled for opportunities to peer at the different food items, we divided the number of peering events directed at each food item by the time that food item was consumed by the association partners on that specific day. We analysed a total of 238 peering events from seven dependent immatures on 25 follow days. In the model, we set the dependent immatures’ peering events (for easy and difficult food items respectively) as the response variable and the time adult association partners spent feeding on easy or difficult food items as the offset term (Model 1). Age of the immatures, processing intensity, and frequency were included as predictors.

#### 2.3.2. Ontogeny of peering

To look at the development of peering over age we used the full data set, i.e., of all immature and adult individuals (see above). We calculated daily peering rates by dividing total peering events by the visible observation duration on each day. The model included peering events as the response variable and the visible observation duration as the offset term (Model 2a). As most peering occurs during immaturity, we focused the more detailed analyses on this age class. To account for different social opportunities to peer, we calculated peering rates that were controlled for the availability of association partners by dividing the number of peering events by the time the immatures spent in peering range to at least one association partner. We included 2993 peering events in this analysis obtained on 538 individual follows. The model included peering events as the response variable, and the time immatures spent in peering range as the offset term (Model 2b).

#### 2.3.3. Development of peering target selection

To investigate the development of the immatures’ peering target selection, we counted and grouped peering events directed at either the mother or non-mother individuals as targets. We used peering events with the mother, and with non-mother targets as response variables, respectively, and visible observation duration as offset term in the models (Models 3a, and 3b). To account for varying opportunities to peer at the two classes of peering targets, we calculated peering rates controlled for the time the observed individual spent in peering range to the respective class of target. To receive proportions of peering, we then divided peering rates by the sum of all peering rates obtained of each class of target. The model included the proportion of immatures’ peering at non-mother targets as the response variable (Model 3c). Further, to investigate the development of peering target selection in terms of peering targets ages’ relative to the age of the peerer, we divided individuals into four age intervals: 0 – 4 years, 4 – 8 years, 8 – 16 years, and 16+ years. Individuals within the same age interval were considered as peers. To account for varying opportunities to peer at the different relative age classes, we calculated proportions of peering as described above. For this analysis we only included data from the wild, and data from Zoo Zürich and Zoo Leipzig as the target selection in Zoo Basel and Zoo Dresden was limited to the immature’s parents due to group composition. This resulted in 238 follows from the wild and 70 follows from the zoos in which peering occurred. In the models, we used the proportions of peering at older, same-aged (‘peers’), and younger individuals as response variables, respectively (Models 3d-f).

#### 2.3.4. Peering context selection

We divided peering contexts into five categories, according to the activity of the peering target: feeding, nesting, social, exploratory, and other (i.e., all non-social activities, such as resting, moving, and grooming of oneself). We calculated the overall proportions of peering in each context by dividing the number of peering events in each context by the total number of peering events for each individual per focal follow day. We also controlled for differing opportunities to peer in each context by first calculating peering rates by dividing the number of peering events in each context by the time adult association partners spent performing the activity per day. We analysed 2797 peering events by 24 dependent immatures (16 from the wild, and 8 from the zoos) obtained on 291 individual follows. We first fitted models with proportions of peering in the respective contexts as the response variable (Models 4a-e). We then compared actual rates of peering, and thus set peering counts in the respective context as the response variable, and the time association partners spent engaging in the behaviour as the offset term (Models 4f-j).

## 3. Results

### 3.1. Peering as a means for social learning in zoo-housed orangutans

Our results revealed significantly higher peering at difficult to process food items compared to easy to process ones (GAMM: Estimate (difficult) = 0.502, p = 0.01; Table 1; Fig. 2a), as well as a significant decrease in peering with increasing frequency of the food item (GAMM: Estimate (Frequency) = - 0.156, p < 0.001; Table 1, Fig. 2b) in zoo-housed orangutans, mirroring previous results on wild orangutans (Schuppli et al. 2016b).

**Figure 1.**
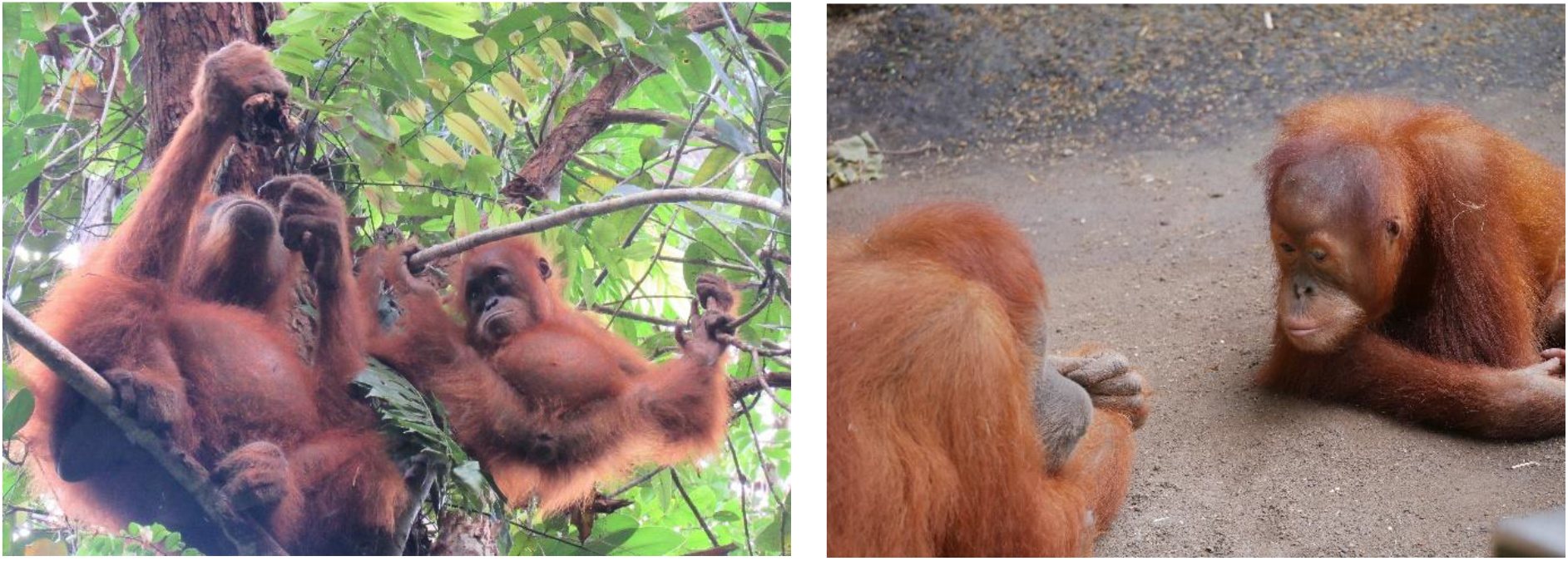
Peering in orangutans. Immature orangutans peering in the feeding context in the wild (left) and in the zoo (right). Photos by Luz Carvajal (left), Ivan Lenzi (right).

**Table 1.** Peering as a measure for social learning: summary of model 1. GAMM with processing intensity of food items (easy / difficult), frequency of food items in association partners’ diet, and age (z-transformed) as fixed effects, and individual and observer ID as random effects. The time adult association partners spent feeding on easy or difficult food items was added as an offset term. Listed are estimates, standard errors, p values, sample size (n), adjusted r^2^, and deviance explained (DE). P values with significance at the 5 % level are indicated in bold font.

| Model | Response variable | Predictors | Type | Estimate | Standard Error | p-value |
| --- | --- | --- | --- | --- | --- | --- |
| 1<br>n = 54<br>$r^2_{adj.} = 0.23$<br>DE = 83.8 % | Peering counts at food items (dependent immatures, zoo only) | <u>Parametric terms:</u> | | | | |
|  |  | (Intercept) | Intercept | 1.544 | 0.877 | 0.078 |
|  |  | Processing intensity (difficult) | Fixed effect | 0.502 | 0.200 | **0.012** |
|  |  | Food item frequency | Fixed effect | -0.156 | 0.028 | **< 0.001** |
|  |  | <u>Smooth terms:</u> |  |  |  |  |
|  |  | Z-Age | Fixed effect |  |  | **< 0.001** |
|  |  | Individual ID | Random effect |  |  | **< 0.001** |
|  |  | Observer ID | Random effect |  |  | 0.585 |

**Figure 2.**
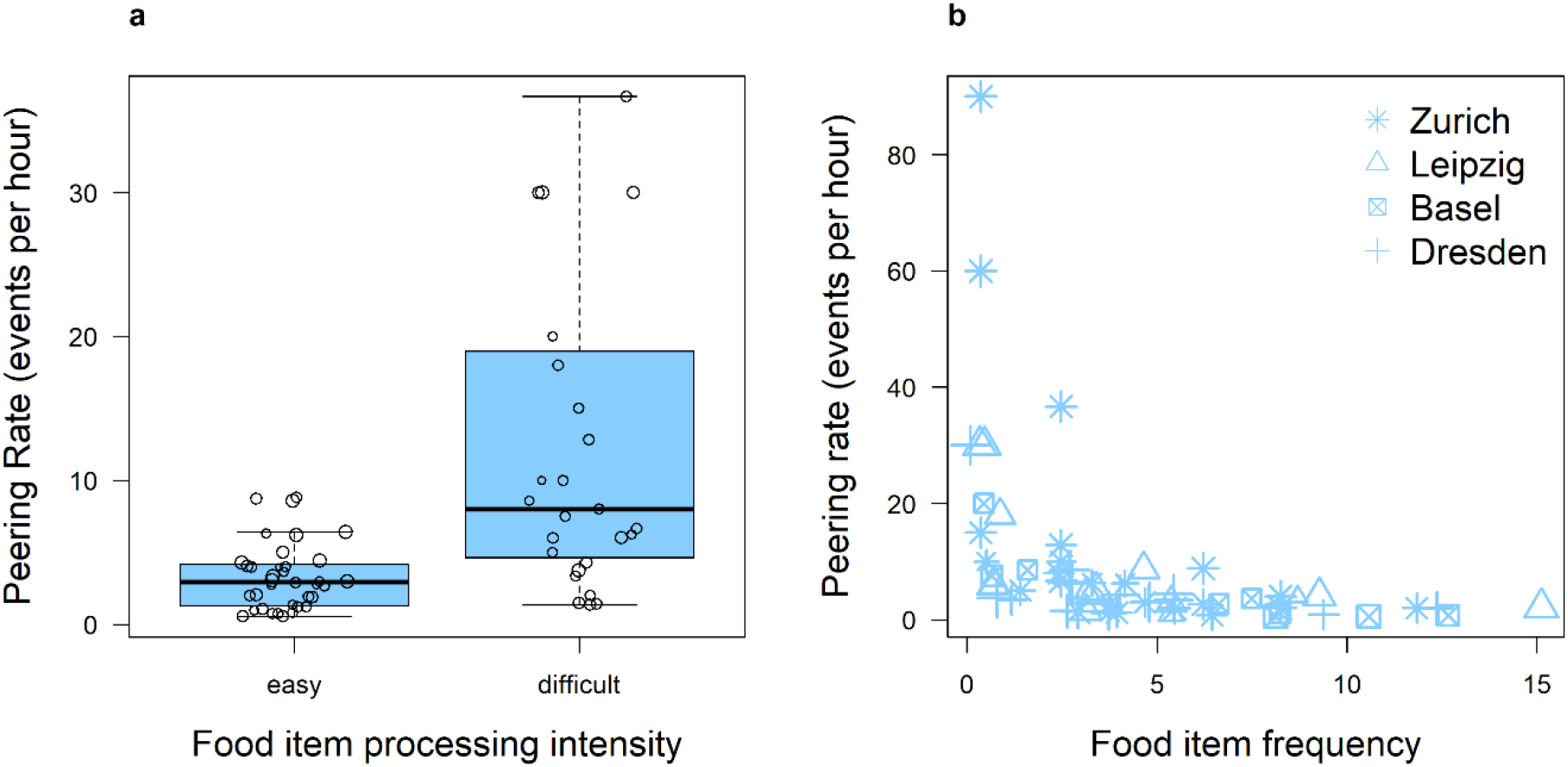
Peering rates in relation to food items’ processing intensity and frequency in association partners’ diet. (a) Mean peering rates (events per hour the food item was consumed) for easy and difficult to process food items of zoo-housed dependent immatures. The bold horizontal line indicates the median, the box the lower and upper quartile, the whiskers the lowest and greatest value, respectively. Empty circles represent the raw data points. (b) Hourly peering rates of zoo-housed dependent immatures over the frequency of food items in adult association partners’ diet. Symbol sizes correspond to the log value of the visible observed hours of each data point.

### 3.2. Ontogeny of peering

In terms of the overall development of peering frequency over the course of an individuals’ life, we found that whereas both settings peering rates showed overall similar non-linear age trajectories, zoo-housed individuals peered significantly more frequently than wild individuals (GAMM: Estimate (Zoo) = 2.395, p < 0.001; Table 2, Model 2a; Fig. 3). A comparison of the age splines for each setting showed significant differences until around the age of six years (Fig. A3): Both in the wild and in the zoos, most peering occurred during the dependency period, however peering rates in the wild peaked around the age of 3 – 4 years, whereas zoo-housed orangutans’ peering rates peaked around the age of 5 – 6 years with a subsequent decrease of peering with age (Fig. 3).

**Table 2.**
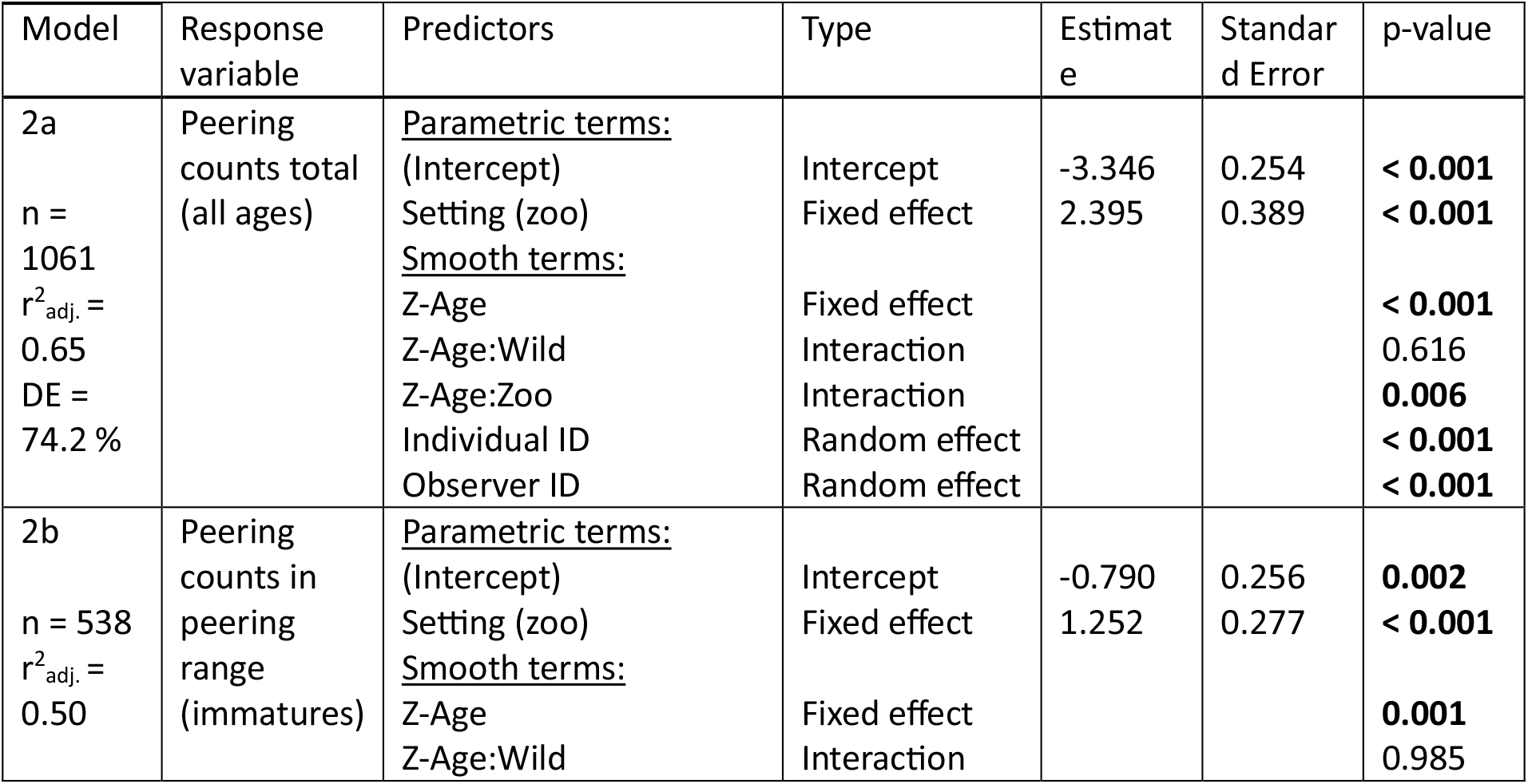
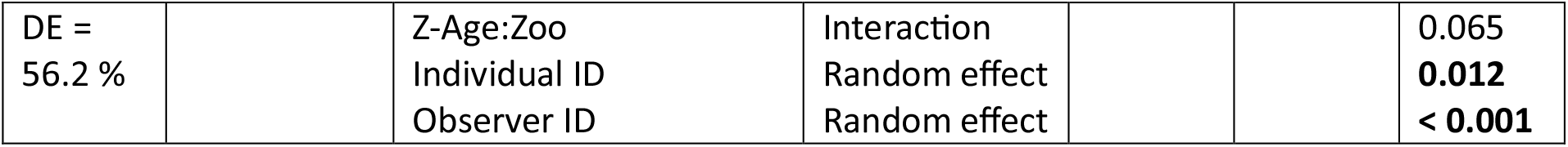
Development of peering frequency over age: summary of models 2a and 2b. GAMMs with setting (Wild / Zoo), age (z-transformed), as well as an interaction between age and setting as fixed effects, and individual and observer ID as random effects. Model 2a included the visible observation duration as the offset term, and model 2b the time immatures spent in peering range. Listed are estimates, standard errors, p values, sample size (n), adjusted r^2^, and deviance explained (DE). P values with significance at the 5 % level are indicated in bold font.

**Figure 3.**
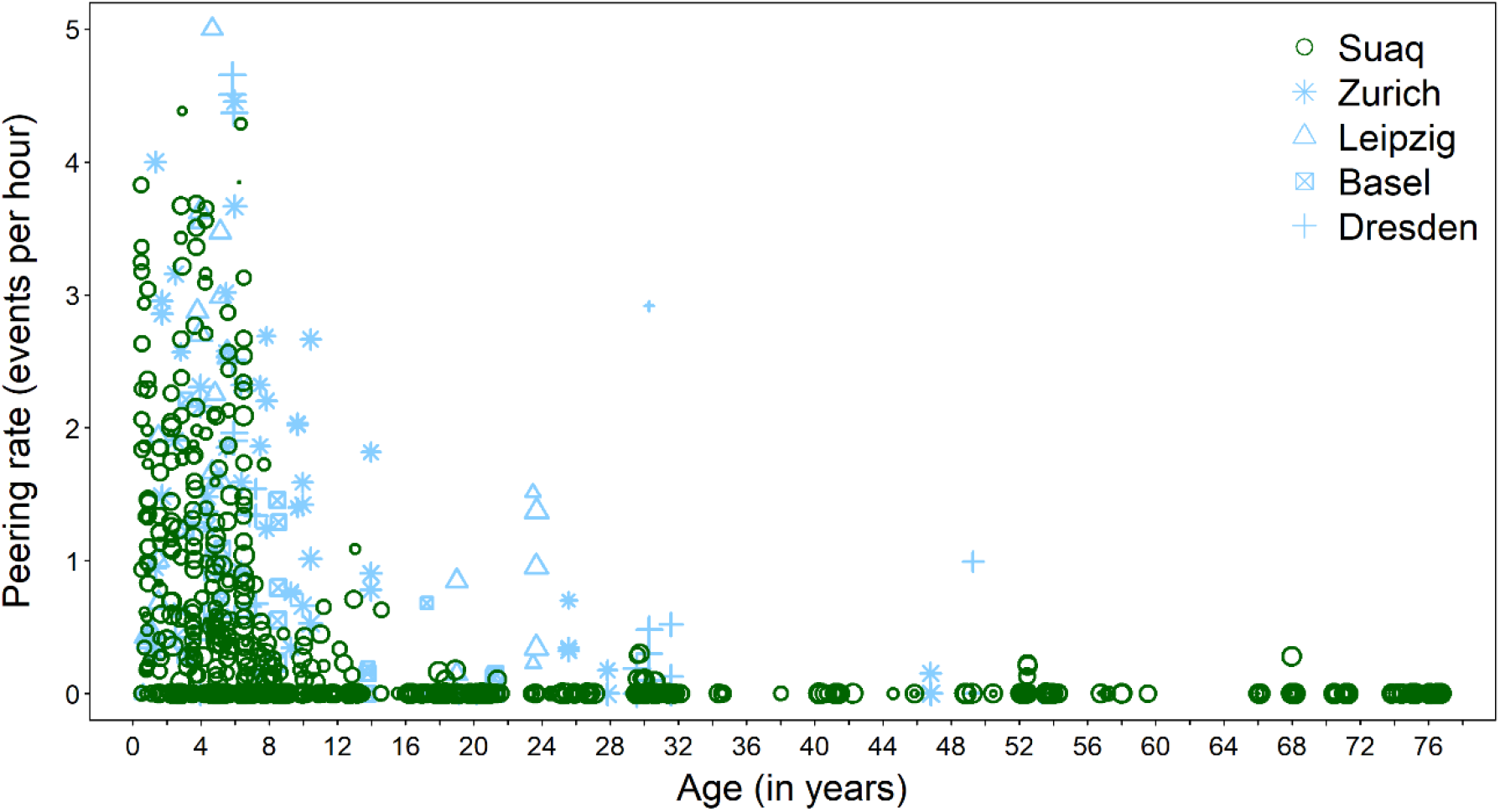
Development of peering frequency. Peering rates (events per visible observation hour) over age in years by wild and zoo-housed orangutans. Each point represents one focal follow day. Symbol sizes correspond to the log value of visible observed hours of each data point.

When looking at peering rates controlled for the availability of association partners within peering range (see methods), we found that zoo-housed immature orangutans peered significantly more frequently than wild immature orangutans (GAMM: Estimate (Zoo) = 1.252, p = < 0.001; Table 2, Model 2b). However, both setting showed overall similar age trajectories of peering rates (Fig. A4, GAMM: p (Age:Wild) = 0.985, p (Age:Zoo) = 0.065; Table 2, Model 2b), with less defined peaks (Fig. 4) compared to overall peering frequencies (i.e., not controlled for the availability of association partners, see Fig. 3).

**Figure 4.**
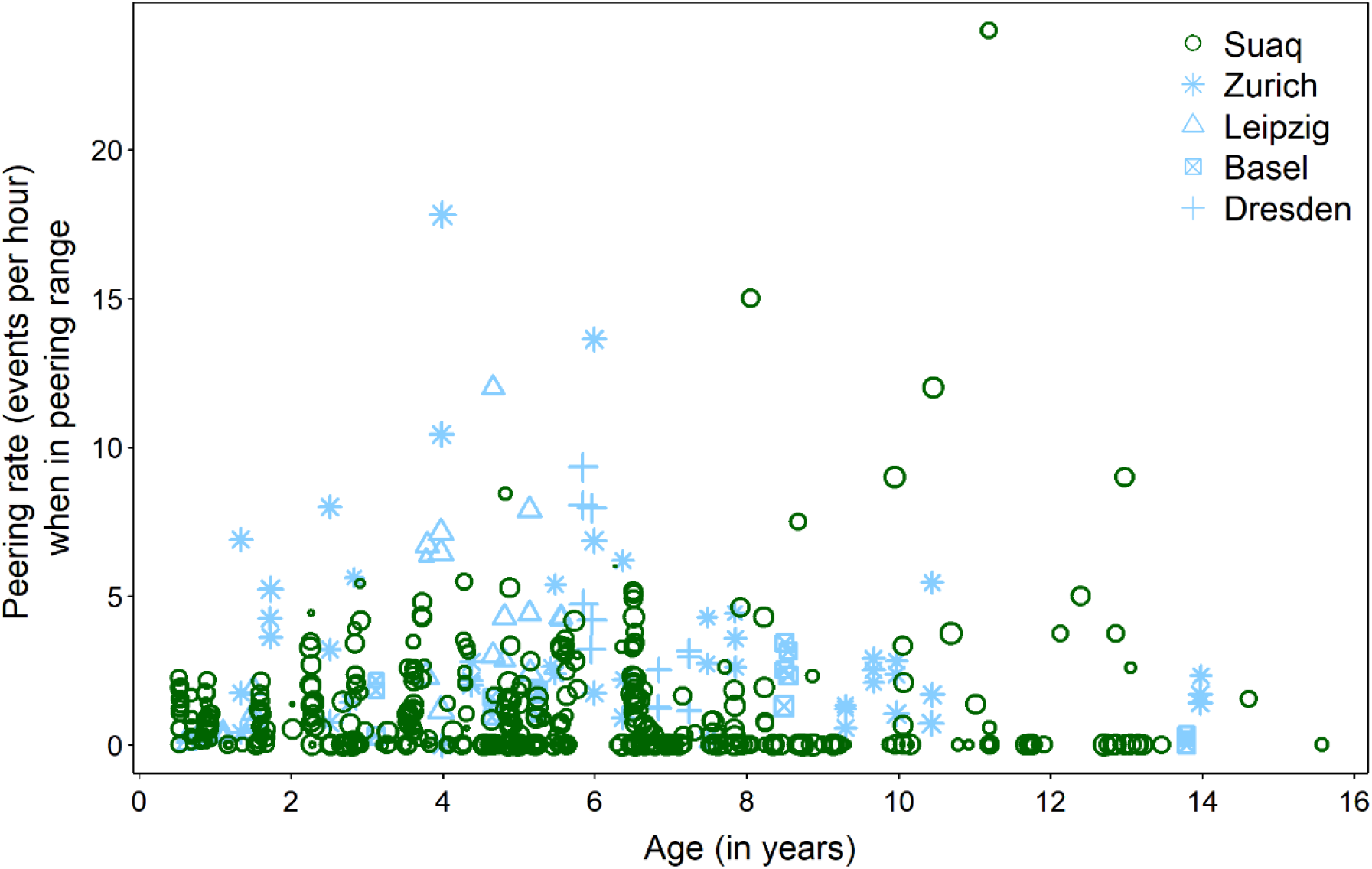
Development of peering frequency controlled for the availability of association partners over age. Peering rates by wild and zoo-housed immature orangutans over age in years calculated as events per hour spent within peering range (0-2 m) of at least one other individual. Each point represents one focal follow day. Symbol sizes correspond to the log value of the visible observed hours of each data point.

### 3.3. Peering target selection

When we compared overall rates of peering directed at the individual’s mother (Fig. 5a) between the settings we found that in both settings, peering at the mother overall decreased over time, but the age trajectories differed between the setting: whereas in the wild, peering rates at the mother decreased linearly with increasing age, in the zoos, the effect of age on peering was non-linear (GAMM: p (Age:Wild) = 0.496, p (Age:Zoo) < 0.001; Table 3, Model 3a). We found that overall rates of peering directed at individuals other than the mother (Fig. 5b) were significantly higher in zoo-housed orangutans compared to wild orangutans (GAMM, Estimate (Zoo) = 3.982, p < 0.001; Table 3, Model 3b), but in neither setting there was a significant age effect (GAMM: p (Age) = 0.659; Table 3, Model 3b).

**Table 3.**
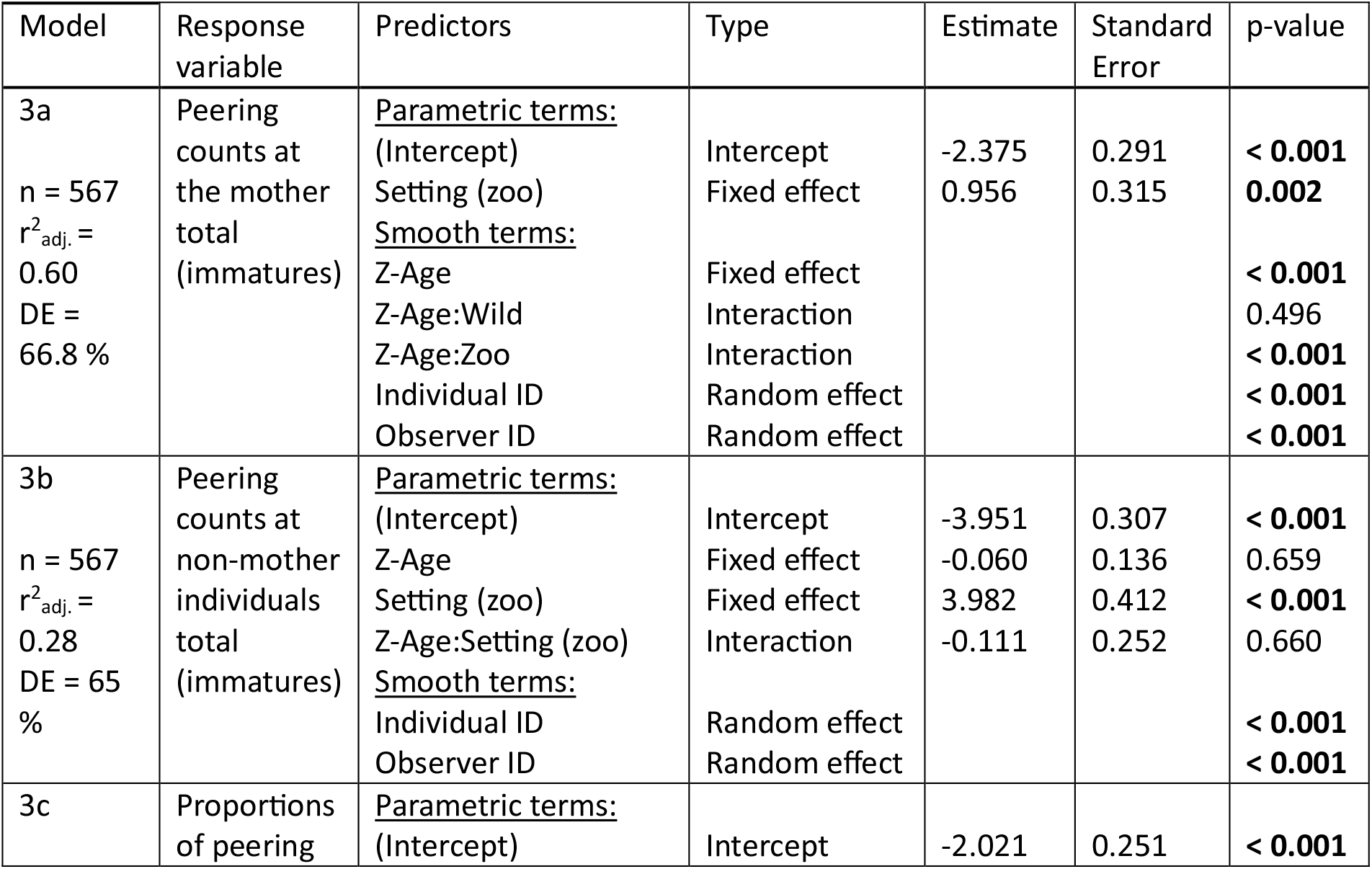
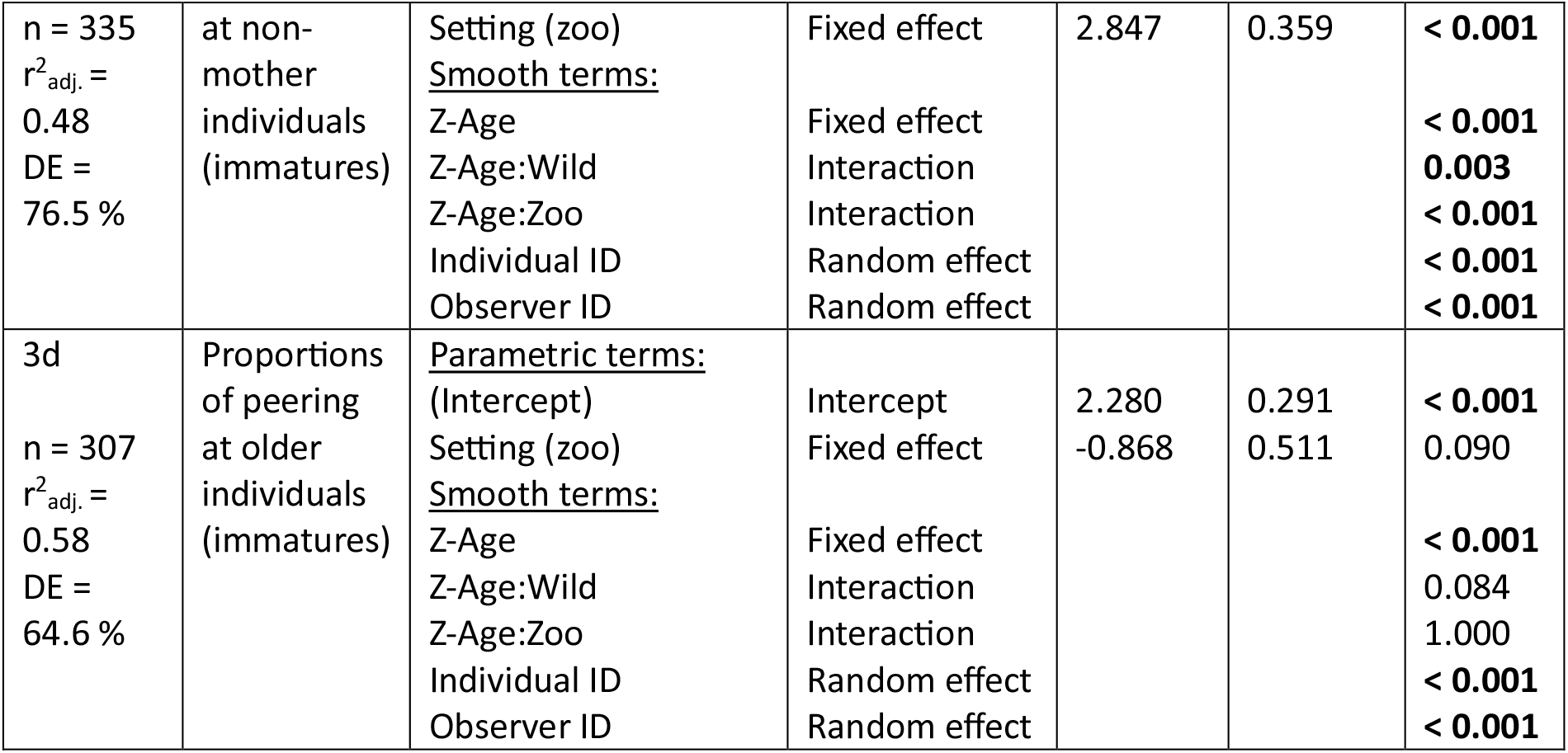
Peering target selection: summary of models 3a-e. GAMMs with setting (Wild / Zoo), age (z-transformed), as well as an interaction between age and setting as fixed effects, and individual and observer ID as random effects. Models 3a and b included the visible observation duration as offset term. Listed are estimates, standard errors, p values, sample size (n), adjusted r^2^, and deviance explained (DE). P values with significance at the 5 % level are indicated in bold font.

**Figure 5.**
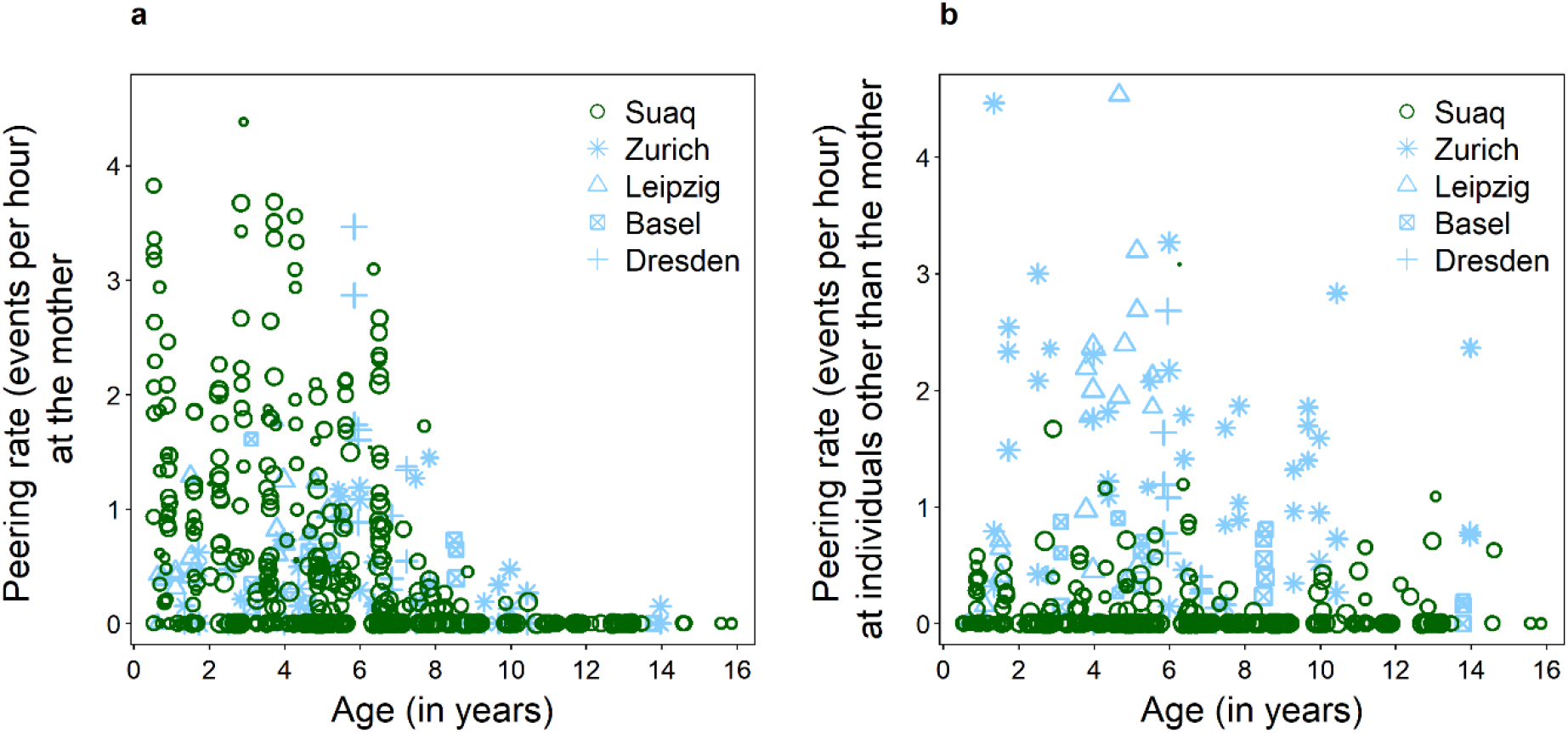
Development of peering frequencies at different peering targets over age. Peering rates (events per visible observation hour) of immature wild and zoo-housed orangutans at (a) their mother, and (b) individuals other than the mother across age in years. Each point represents one focal follow day. Symbol sizes correspond to the log value of the visible observed hours of each data point.

When we controlled for varying opportunities to peer at the two different target classes (see methods), we found that zoo-housed immature orangutans directed a significantly larger proportion of their peering at non-mother individuals than wild immature orangutans (GAMM: Estimate (Zoo) = 2.847, p < 0.001; Table 3, Model 3c, Fig. 6a). Age had a non-linear effect on these peering proportions (GAMM: p (Age) < 0.001, Table 3, Model 3c, Fig. 6a). Furthermore, the age patterns differed between wild and zoo-housed immatures (Fig. A6): In the wild, peering at non-mother individuals increased over time, whereas this clear shift was absent in the zoos (Fig. 6a).

**Figure 6.**
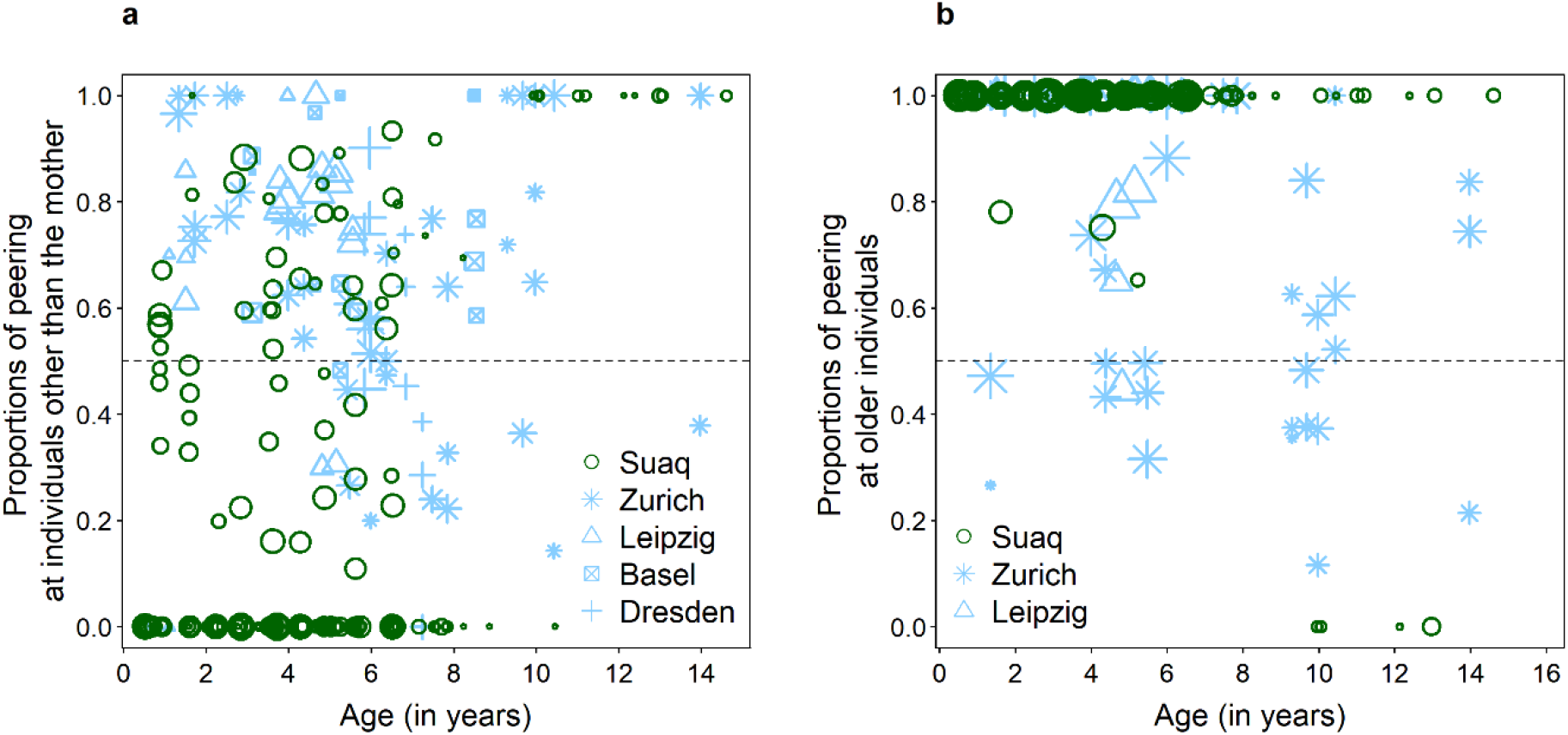
Development of peering frequencies at different peering targets over age controlled for opportunities to peer: (a) mother vs. non-mother individuals, and (b) older peering targets. Proportions of peering by immatures directed at (a) non-mother individuals in the wild and in the zoo, and (b) older individuals across age in years controlled for the time the peering individual spent in close proximity to each class of peering target. Each point represents one focal follow day. The size of the symbols reflects the log value of the total number of observed peering events for each data point. The dashed line at 0.5 indicates an equal proportion of peering at the mother and other individuals.

When looking at peering at targets of different ages in relation to the peerer’s age (controlled for varying opportunities to peer at the different age classes, see methods), we found no significant differences in peering proportions between wild and zoo-housed immatures (Fig. 6b and A7, Table 3, Models 3d, and Table A5, Models 3d, e). Both wild and zoo-housed orangutans directed most of their peering at older and a very low share of their peering at younger individuals (Fig. A7b).

### 3.4. Peering context selection

We found that in both settings, overall, most peering occurred in the feeding context (82.3 % in the wild, 49.9 % in the zoos). Wild orangutans directed about 13.7 % of peering at nest-building behaviour, but hardly peered at social (1.0 %) or exploratory behaviour (1.2 %). In contrast, zoo-housed immature orangutans frequently peered in exploratory (34.0 %) or social (7.8 %) contexts, but only directed about 1.7 % of peering at nesting activities (Fig. 7a). These differences in peering proportions were significant for all contexts (Fig. 7a; see supplementary material for model summaries: Table A6, Models A4a-e). Note that the response variables in the five models on peering proportions (Table A6, Models A4a-e) were not independent of each other, however, the p-values of the models remain significant even after correcting for multiple comparisons.

**Figure 7.**
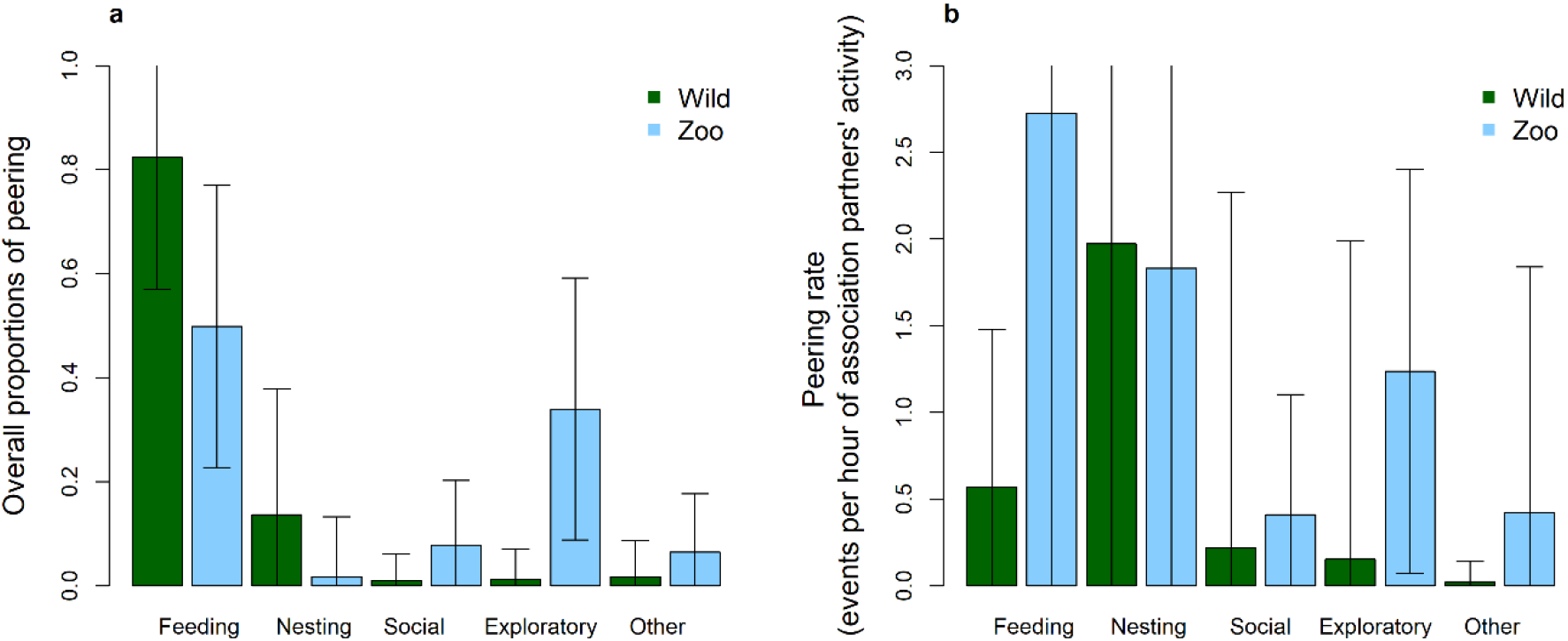
Contexts in which peering occurred. Peering by wild and zoo-housed dependent immature orangutans directed at different activities of their association partners, namely feeding, nesting, social interactions, exploratory behaviour and other, non-social activities. (a) shows overall proportions of peering, and (b) rates of peering controlled for the time association partners spent engaging in the respective activity. The bars represent the mean proportions (a) and rates (b) per focal follow day. Arrows represent the standard deviations, note that the top range of some standard deviation is cut off.

When looking at rates of peering controlled for the time association partners engaged in the activity (see methods), we found that wild immatures most frequently peered at activities in the feeding and nest-building contexts, followed by social activities (Fig. 7b). Zoo-housed immatures peered most frequently in the feeding context as well as the nest-building context, followed by explorative contexts (Fig. 7b). These differences between wild and zoo-housed immatures were not significant, with the exception of the feeding context where zoo-housed immatures peered significantly more (GAMM: Estimate (Zoo) = 1.560, p < 0.001; Table A6, Model A6f).

## 4. Discussion

In this study we compared peering behaviour of wild and zoo-housed Sumatran orangutans to investigate how this behaviour is regulated on the immediate and developmental level, to ultimately shed light on the mechanisms underlying orangutan observational social learning. Specifically, we aimed to disentangle the extent to which peering is influenced by immediate and developmental social and ecological factors, as opposed to being hard-wired (i.e., not affected by immediate conditions (Bouchard et al. 2007) or conditions that the individual had experienced during development and thus at least to some extent genetically determined). To that end, we compared peering rates across age, peering targets, and contexts between the two settings.

### 4.1. Peering as a means for social learning

Attending to social information via observation is a cornerstone of observational social learning, whereas the difference between the observed information and the existing knowledge of the peering individual determines whether learning takes place. Wild immature orangutans have been found to selectively peer in learning intense contexts, such as rare and difficult to process food items (Schuppli et al. 2016b). As predicted, we found that zoo-housed dependent immatures peered significantly more at difficult to process food items, and for items that are rare in their diets (Fig. 2, Table 1). These findings support that both wild and in zoos immature orangutans use peering to learn in the foraging context.

### 4.2. Ontogeny of peering

We found that peering rates in both wild and zoo-housed orangutans followed similar age trajectories, with most peering occurring during the dependency period (Fig. 3), suggesting that the tendency to peer is independent of immediate or developmental social or ecological factors and thus likely follows a hard-wired developmental trajectory. However, overall peering rates were higher in zoo-housed orangutans, indicating that either immediate or developmental factors influence the frequency at which the behaviour is expressed (Fig. 3, Table 2, Model 2a). Peering rates in the wild peaked slightly earlier, around 3 – 4 years of age, compared to 5 – 6 years in zoo-housed orangutans. This is surprising, considering that zoo-housed orangutans tend to show shorter developmental periods compared to their conspecifics in the wild which suggests an overall faster physical development (Markham 1995; Shumaker et al. 2008; van Noordwijk et al. 2018). Likely, increased social opportunities to peer at targets other than the mother, as well as more available free time (Dalimunthe et al. 2021) enable zoo-housed orangutans to peer at a higher frequency, for a drawn-out time during their development. Even after we controlled for differences in social opportunities, zoo-housed immature orangutans still exhibited higher rates of peering compared to wild immatures (Fig. 4, Table 2, Model 2b), suggesting that peering frequency is not solely a consequence of social opportunities to peer.

Peering requires that the peering target tolerates the peering individual in close spatial proximity. Schuppli et al. (2017) found that in the wild, immatures of the more sociable and tolerant Sumatran orangutan species show significantly higher peering rates compared to the less social Bornean orangutans. While the 2017 study could not differentiate between developmental and intrinsic effects on peering rates, the results of our study on the same species suggest that differences in peering rates are to some extent caused by developmental factors but that immediate ecological factors also play a role (see 4.3 below). For wild immature orangutans, their own mother was the main peering target, and thus her levels of tolerance may shape peering opportunities. In the wild, a mother’s tolerance levels regarding food interactions peak during the time her offspring learns feeding skills, which occurs around 3 – 5 years (Mikeliban et al. 2021), coinciding with the peak in peering rates observed here. After that, tolerance by the mother gradually decreases, most likely because she starts to energetically prepare for a new pregnancy (van Noordwijk 2012; Falkner 2015). Because they are provisioned, zoo-housed mothers likely do not experience these energetic constraints to the same extent as mothers in the wild and may therefore maintain high levels of tolerance throughout their offspring’s dependency period. Further, zoo-housed immature orangutans are in constant association with other individuals aside from their mother, and thus peering opportunities are not dependent on her tolerance alone. In the wild, interaction with the mother tends to decrease with increasing group size, suggesting that social dynamics influence immature orangutan behaviour (Fröhlich et al. 2020).

### 4.3. Development of peering target selection

In both wild and zoo-housed immature orangutans, peering directed at the mother decreased with increasing age of the individual (Fig. 5a), which implies a hard-wired predisposition to peer at the mother during dependency. The higher peering rates at the mother in zoo-housed immatures were likely a consequence of their overall higher frequency of peering (see above). Peering rates at non-mother individuals were also higher in zoo-housed immatures compared to wild immatures whereby the difference was more pronounced than for peering at the mother (Table 3, Model 3b). Furthermore, peering at individuals other than the mother increased with age in the wild, whereas there was no discernible pattern in zoo-housed immature orangutans (Fig. 5b). This suggests that social and ecological factors influence the more nuanced patterns of peering target selection across different ages. When controlling for social opportunities, immature zoo orangutans exhibited a higher overall preference to peer at non-mother individuals compared to wild immatures (Fig. 6a). In the wild, peering at non-mother individuals gradually increases with age and after the age of ten individuals almost exclusively peer at them (Schuppli et al. 2016b; Schuppli and van Schaik 2019a), coinciding with increased ranging independence from their mothers (van Noordwijk et al. 2009). However, this age-specific pattern was absent in zoo-housed orangutans. Wild orangutans show a strong tendency to peer at relatives (Schuppli and van Schaik 2019a), and tolerance towards kin is usually higher than towards unrelated individuals (Pisor and Surbeck 2019). Most of the orangutans from our zoo populations were either to some degree related or spent most of their lives together. Growing up closely surrounded by the same individuals may establish group members other than the mother as trusted role models for the immatures which may lead to higher peering rates. Further, a recent study found evidence that favourable environmental conditions, such as food availability, positively affect social learning opportunities and tolerance between association partners (Mörchen et al. 2024). Unlike in the wild, zoo-housed orangutans are regularly provisioned with food and experience little to no resource competition.

All in all, these findings are consistent with the observation of higher peering frequencies at the mother in Sumatran orangutans compared to Bornean orangutans for the feeding context, even after controlling for varying opportunities to peer and food item processing intensity (Schuppli et al. 2017). The authors attributed these findings both to potential innate differences between the two species as well as potential developmental effects of either increased sociality, or broader skill sets in the population and thus more for the immature to learn. In combination with the results of the current study, it appears likely that sociality and tolerance of a population experienced during development influence peering behaviour in immatures, and that immediate factors, particularly the availability of food sources (which affects an individual’s energetic state and the time that it can devote to activities other than feeding), may further influence the frequency of the behaviour.

We found that immatures in both settings preferentially peered at older individuals or at peers but rarely at younger conspecifics (Fig. 6b, and A7). This is in line with previous findings on selective bias in chimpanzees (Biro et al. 2003) and capuchin monkeys (Barrett et al. 2017; Coelho et al. 2015). However, zoo-housed orangutans receive less variable food which usually requires little pre-ingestive processing (see Fig. A8 for a comparison of the diet processing intensity of the two settings). Consequently, individuals growing up in zoos are likely to possess complete skill sets at an earlier age compared to their wild conspecifics, which makes it less likely that peering individuals will indeed learn more by peering at older individuals compared to younger individuals or peers. The fact that we nevertheless find a bias to peer at older individuals in zoo-housed orangutans suggests a hard-wired bias to select older individuals. A hard-wired tendency to peer at older individuals is advantageous for species adapted to learning-intense foraging niches, because in natural settings, older individuals are likely to be experts and learning from them is more efficient and safer (Henrich and Broesch 2011).

### 4.4. Peering context selection

Wild and zoo-housed immature orangutans differed in the contexts in which they peered (Figure 7a, Table A6, Models A6a-e). When looking at absolute peering proportions, immatures in the zoos spent less time peering in feeding and nest-building contexts compared to wild orangutans, and more time peering at social and exploratory behaviours. Nest-building in the wild is an intricate and multistep process which includes bending and intertwining branches, as well as adding layers of pillows and blankets made from leavy twigs and loose leaves (Prasetyo et al. 2009). In zoos, nest-building mainly consists of gathering wood wool, blankets, or other soft materials provided for the animals. Similarly, feeding skills are less processing intense in zoos (Fig. A8). Consequently, zoo-housed orangutans may reach adult-like levels of proficiency in these skills earlier than their wild conspecifics, leaving them with more time to direct their attention to other activities. Similarly, wild Bornean orangutans whose foraging techniques contain fewer steps of pre-ingestive processing compared to those of Sumatran orangutans, show lower proportions of peering in the feeding context (Schuppli et al. 2016b). Exploratory and social behaviours are generally rare in wild orangutans (Morrogh-Bernard et al. 2009; van Schaik et al. 2016), due to their strong neophobia and solitary lifestyle. In wild dependent immatures, opportunities to peer in social contexts are restricted to when their mothers are associating with other individuals (Lonsdorf 2006). Indeed, relative to the time association partners spent engaging in the respective activity (i.e., controlled for the opportunities to peer in these different contexts), wild immature orangutans peered most in nest-building, feeding, and social contexts (Figure 7b, see also Schuppli et al. 2016b), and zoo-housed immatures mostly peered in feeding, nest-building and exploratory contexts (Fig. 7b). However, these differences in peering rates controlled for peering opportunities between wild and zoo-housed immatures were not significant, with the exception of the feeding context. These results suggest that immatures selectively peer in contexts that are either rare or more learning intense, given their ecological and social environment.

## 5. Conclusion

Our findings suggest that an immature orangutan’s general tendency to peer is independent of their social and ecological environment and thus likely to some extent hard-wired. In other words, the drive to attend to social information seems to develop largely independent of experience and seems mostly unaffected by prevailing conditions. However, our findings further show that the frequency at which the behaviour is expressed as well as its more fine-grained patterns are likely influenced by immediate and developmental social and ecological factors. Among other potential factors, social complexity, social tolerance and food availability during development seem to modulate peering behaviour. Given that peering is a measure of observational social learning, we can thus conclude that orangutan observational social learning has a genetically fixed foundation but that it also entails a large degree of behavioural plasticity allowing individuals to adjust to the specific social and ecological conditions during their lives. Therefore, even though orangutans are the least sociable of the great apes (Goossens et al. 2009), they seem to be capable of complex social behaviours when different conditions demand it (Edwards and Snowdon 1980). Further, the ability to flexibly adjust learning behaviour in response to changing social and ecological conditions has likely contributed to the development of the vast repertoires and rich culture great apes possess today (Whiten et al. 1999; van Schaik and Burkart 2011), and may have also been a key factor in the evolution of human cognition (Boyd et al. 2011).

Lastly, this study highlights the value of comparative studies of captive and wild populations of the same species. Our results exemplify that comparing the same species under a large range of social and ecological conditions can give us a more complete picture of a behaviour and shed light on the underlying mechanisms at play.

## Acknowledgements

We thank all researchers, and students who conducted projects on peering at the Suaq Balimbing research site, and whose data were essential for this study, namely Natasha Bartolotta, Helvi Musdarlia, Anais van Cauwenberghe, Tri Rahmaeti, Sonja Falkner, Luz Carvajal, Ellen Meulmann, Sofia Forss, Olivia Wassmer, Belinda Kunz, Beatrice Ehmann, Andrea Permana, and Julia Mörchen. We acknowledge all students, volunteers, and local field assistants involved in the collection of standard behavioural data for the long-term database of Suaq Balimbing. We are thankful to the technicians, and students who collected standard behavioural data in the zoos, especially to Ivan Lenzi, Francois Lamarque, Sree Subha Ramaswamy, Glenn Honstetter, Anais van Cauwenberghe, Shubhangi Kansal, Nele Käter, and Mulati Mikeliban. Further, we gratefully acknowledge the curators, staff, and helpers at the zoo for their support to conduct our data collection: Adrian Baumeyer (Zoo Basel), Dr. Claudia Rudolf von Rohr (Zoo Zurich), Dr. Leyla Davis (Zoo Zurich), Dr. Wolfgang Ludwig (Zoo Dresden), Roman Richter (Zoo Dresden). We thank our colleagues at the Max Planck-Institute for Evolutionary Anthropology in Leipzig, and especially Hanna Petschauer, and Dr. Daniel Hanus. We gratefully acknowledge the Badan Riset dan Inovasi Nasional (BRIN), the Sumatran Orangutan Conservation Survival Foundation (SOCP), the Universitas Nasional (UNAS), as well as the Taman Nasional Gunung Leuser (TNGL) for their permission and support for this research.

## Ethics approval and consent to participate

This study on wild and zoo-housed orangutans was strictly observational and non-invasive, and there was no interaction with our study animals. The research protocols for the observations at Suaq Balimbing were approved by the Indonesian State Ministry for Research, Technology and Higher Education (RISTEK; Research Permit No.: 152/SIP/FRP/SM/V/2012 and following) and complied with the legal requirements of Indonesia. At the zoos, the orangutans were observed from the visitor areas (or in the case of Leipzig Zoo, an observation tower). The observations did not interfere with the daily routine of the orangutans. The study complies with the Weatherfall report titled ‘The use of non-human primates in research’ (Academy of Medical Sciences & Weatherall, 2006), the EAZA Minimum Standards for the Accommodation and Care of Animals in Zoos and Aquaria (EAZA, 2014), the WAZA Ethical Guidelines for the Conduct of Research on Animals by Zoos and Aquariums (Mellor et al., 2015), and the ASAB/ABS’s Guidelines for the Treatment of Animals in Behavioural Research and Teaching (Buchanan et al., 2012). IAUCUC approval was not necessary to conduct this research. A joint ethical committee of the Max Planck Institute for Evolutionary Anthropology and Leipzig Zoo approved this study.

## Data availability statement

The data set for this article will be deposited to an online repository *upon publication*.

## References

Afseth, Cassandra; Shim, Andrew; Anderson, Samantha; Bell, Alison M.; Hellmann, Jennifer K. (2022): Vertical transmission of horizontally acquired social information in sticklebacks: implications for transgenerational plasticity. In Proceedings. Biological sciences 289 (1979), p. 20220571. DOI: 10.1098/rspb.2022.0571.

Barrett, Brendan J.; McElreath, Richard L.; Perry, Susan E. (2017): Payoff-biased social learning underlies the diffusion of novel extractive foraging traditions in a wild primate.

Benson-Amram, Sarah; Weldele, Mary L.; Holekamp, Kay E. (2013): A comparison of innovative problem-solving abilities between wild and captive spotted hyaenas, Crocuta crocuta. In Animal Behaviour 85 (2), pp. 349–356. DOI: 10.1016/j.anbehav.2012.11.003.

Biro, Dora; Inoue-Nakamura, Noriko; Tonooka, Rikako; Yamakoshi, Gen; Sousa, Claudia; Matsuzawa, Tetsuro (2003): Cultural innovation and transmission of tool use in wild chimpanzees: evidence from field experiments. In Animal cognition 6 (4), pp. 213–223. DOI: 10.1007/s10071-003-0183-x.

Boesch, Christophe (2021): Identifying animal complex cognition requires natural complexity. In iScience 24 (3), p. 102195. DOI: 10.1016/j.isci.2021.102195.

Bouchard, Julie; Goodyer, William; Lefebvre, Louis (2007): Social learning and innovation are positively correlated in pigeons (Columba livia). In Animal cognition 10 (2), pp. 259–266. DOI: 10.1007/s10071-006-0064-1.

Boyd, Robert; Richerson, Peter J.; Henrich, Joseph (2011): The cultural niche: why social learning is essential for human adaptation. In Proceedings of the National Academy of Sciences of the United States of America 108 Suppl 2 (Suppl 2), pp. 10918–10925. DOI: 10.1073/pnas.1100290108.

Carvajal, Luz; Schuppli, Caroline (2022): Learning and skill development in wild primates: toward a better understanding of cognitive evolution. In Current Opinion in Behavioral Sciences 46, p. 101155. DOI: 10.1016/j.cobeha.2022.101155.

Coelho, C. G.; Falótico, T.; Izar, P.; Mannu, M.; Resende, B. D.; Siqueira, J. O.; Ottoni, E. B. (2015): Social learning strategies for nut-cracking by tufted capuchin monkeys (Sapajus spp.). In Animal cognition 18 (4), pp. 911–919. DOI: 10.1007/s10071-015-0861-5.

Dalimunthe, Nurzaidah Putri; Alikodra, Hadi Sukadi; Iskandar, Entang; Utami-Atmoko, Sri Suci (2021): The activity budgets of captive orangutan (Pongo pygmaeus) in two different Indonesian zoos. In Biodiversitas 22 (4). DOI: 10.13057/biodiv/d220438.

Damerius, Laura A.; Forss, Sofia I. F.; Kosonen, Zaida K.; Willems, Erik P.; Burkart, Judith M.; Call, Josep et al. (2017): Orientation toward humans predicts cognitive performance in orang-utans. In Scientific reports 7, p. 40052. DOI: 10.1038/srep40052.

Edwards, Sara D.; Snowdon, Charles T. (1980): Social behaviour of captive, group-living Orangutans. In International Journal of Primatology 1 (1), pp. 39–62. DOI: 10.1007/BF02692257.

Falkner, Sonja (2015): Mother-Offspring Conflict in Orangutans. Disentangling different contexts of mother-offspring conflict in Sumatran and Bornean orangutans. Masters. University of Zurich, Zurich. Anthropological Institute and Museum.

Forss, Sofia I. F.; Schuppli, Caroline; Haiden, Dominique; Zweifel, Nicole; van Schaik, Carel P. (2015): Contrasting responses to novelty by wild and captive orangutans. In American Journal of Primatology 77, pp. 1109–1121.

Forss, Sofia Ingrid Fredrika; Motes-Rodrigo, Alba; Dongre, Pooja; Mohr, Tecla; van de Waal, Erica (2022): Captivity and habituation to humans raise curiosity in vervet monkeys. In Animal cognition 25 (3), pp. 671–682. DOI: 10.1007/s10071-021-01589-y.

Fröhlich, Marlen; Kunz, Julia; Fryns, Caroline; Falkner, Sonja; Rukmana, Evasari; Schuppli, Mélanie et al. (2020): Social interactions and interaction partners in infant orang-utans of two wild populations. In Animal Behaviour 166, pp. 183–191. DOI: 10.1016/j.anbehav.2020.06.008.

Goossens, Benoît; Chikhi, Lounès; Fairus Jalil, Mohd.; James, Sheena; Ancrenaz, Marc; Lackman-Ancrenaz, Isabelle; Bruford, Michael W. (2009): Taxonomy, geographic variation and population genetics of Bornean and Sumatran orangutans. In Serge A. Wich, S. Suci Utami Atmoko, Tatang Mitra Setia, Carel P. van Schaik (Eds.): Orangutans: Geographic Variation in Behavioral Ecology and Conservation: Oxford University Press, pp. 1–14.

Grampp, Mathilde; Sueur, Cédric; van de Waal, Erica; Botting, Jennifer (2019): Social attention biases in juvenile wild vervet monkeys: implications for socialisation and social learning processes. In Primates; journal of primatology 60 (3), pp. 261–275. DOI: 10.1007/s10329-019-00721-4.

Harrison, Rachel A.; van de Waal, Erica (2022): The unique potential of field research to understand primate social learning and cognition. In Current Opinion in Behavioral Sciences 45, p. 101132. DOI: 10.1016/j.cobeha.2022.101132.

Haslam, Michael (2013): ‘Captivity bias’ in animal tool use and its implications for the evolution of hominin technology. In Philosophical transactions of the Royal Society of London. Series B, Biological sciences 368 (1630), p. 20120421. DOI: 10.1098/rstb.2012.0421.

Henrich, Joseph; Broesch, James (2011): On the nature of cultural transmission networks: evidence from Fijian villages for adaptive learning biases. In Philosophical transactions of the Royal Society of London. Series B, Biological sciences 366 (1567), pp. 1139–1148. DOI: 10.1098/rstb.2010.0323.

Herrmann, Esther; Call, Josep; Hernàndez-Lloreda, Maráa Victoria; Hare, Brian; Tomasello, Michael (2007): Humans have evolved specialized skills of social cognition: the cultural intelligence hypothesis. In Science (New York, N.Y.) 317 (5843), pp. 1360–1366. DOI: 10.1126/science.1146282.

Heyes, C. M. (1994): Social learning in animals: categories and mechanisms. In Biological Reviews 69, pp. 207–231. DOI: 10.1111/j.1469-185X.1994.tb01506.x.

Heyes, Cecilia (2012): What’s social about social learning? In Journal of comparative psychology (Washington, D.C.: 1983) 126 (2), pp. 193–202. DOI: 10.1037/a0025180.

Hoppitt, William; Laland, Kevin N. (2013): Social Learning. An Introduction to Mechanisms, Methods, and Models: Princeton University Press. Available online at http://www.jstor.org/stable/j.ctt2jc8mh.

Idani, Gen’ichi (1995): Function of peering behaviour among bonobos (Pan paniscus) at Wamba, Zaire. In Primates 36 (3), pp. 377–383.

Jaeggi, Adrian V.; Dunkel, Lynda P.; van Noordwijk, Maria A., Wich Serge A.; Sura, Agnes A.L.; van Schaik, Carel (2010): Social learning of diet and foraging skills by wild immature Bornean orangutans: Implications for culture. In American Journal of Primatology 72, pp. 62–71.

Jaeggi, Adrian V.; van Noordwijk, Maria; van Schaik, Carel P. (2008): Begging for information: mother– offspring food sharing among wild Bornean orangutans. In American Journal of Primatology (70), pp. 533–541.

Johnson, Christine M.; Frank, Rebecca E.; Flynn, Danielle (1999): Peering in mature, captive bonobos (Pan paniscus). In Primates 40 (2), pp. 397–407.

Kummer, Hans; Goodall, Jane (1985): Conditions of innovative behaviour in primates. In Philosophical Transactions of the Royal Society of London Series B: Biological Sciences 308 (1135), pp. 203–214. DOI: 10.1098/rstb.1985.0020.

Lenth, Russel V. (2023): emmeans: Estimated Marginal Means, aka Least-Squares Means. Available online at https://CRAN.R-project.org/package=emmeans.

Lonsdorf, Elizabeth V. (2006): What is the role of mothers in the acquisition of termite-fishing behaviors in wild chimpanzees (Pan troglodytes schweinfurthii)? In Animal cognition 9 (1), pp. 36–46. DOI: 10.1007/s10071-005-0002-7.

Markham, R. (1995): Doing It Naturally. In Ronald D. Nadler, Birute F. M. Galdikas, Lori K. Sheeran, Norm Rosen (Eds.): The Neglected Ape. Boston, MA: Springer US, pp. 273–278.

Matsuzawa, Tetsuro; Biro, Dora; Humle, Tatyana; Inoue-Nakamura, Noriko; Tonooka, Rikako; Yamakoshi, Gen (2001): Emergence of Culture in Wild Chimpanzees: Education by Master-Apprenticeship. In Tetsuro Matsuzawa (Ed.): Primate Origins of Human Cognition and Behavior. Tokyo: Springer Japan, pp. 557–574.

Mendonça, Renata S.; Kanamori, Tomoko; Kuze, Noko; Hayashi, Misato; Bernard, Henry; Matsuzawa, Tetsuro (2017): Development and behaviour of wild infant-juvenile East Bornean orangutans (Pongo pygmaeus morio) in Danum Valley. In Primates; journal of primatology 58 (1), pp. 211–224. DOI: 10.1007/s10329-016-0567-6.

Mikeliban, Mulati; Kunz, Belinda; Rahmaeti, Tri; Uomini, Natalie; Schuppli, Caroline (2021): Orangutan mothers adjust their behaviour during food solicitations in a way that likely facilitates feeding skill acquisition in their offspring. In Scientific reports 11 (1), p. 23679. DOI: 10.1038/s41598-021-02901-z.

Mitra Setia, T.; Delgado, R. A.; Utami Atmoko, S. S.; Singleton, I.; van Schaik, C. P. (2009): Social organization and male-female relationships. In Orangutans: Geographic Variation in Behavioral Ecology and Conservation. DOI: 10.5167/UZH-31343.

Mörchen, Julia; Luhn, Frances; Wassmer, Olivia; Kunz, Julia A.; Kulik, Lars; van Noordwijk, Maria A. et al. (2023): Migrant orangutan males use social learning to adapt to new habitat after dispersal. In Front. Ecol. Evol. 11, Article 1158887. DOI: 10.3389/fevo.2023.1158887.

Mörchen, Julia; Luhn, Frances; Wassmer, Olivia; Kunz, Julia A.; Kulik, Lars; van Noordwijk, Maria A. et al. (2024): Orangutan males make increased use of social learning opportunities, when resource availability is high. In iScience 27 (2), p. 108940. DOI: 10.1016/j.isci.2024.108940.

Morrogh-Bernard, Helen C.; Husson, Simon J.; Knott, Cheryl D.; Wich, Serge A.; van Schaik, Carel P.; van Noordwijk, Maria A. et al. (2009): Orangutan activity budgets and diet: A comparison between species, populations and habitats. In Serge A. Wich, S. Suci Utami Atmoko, Tatang Mitra Setia, Carel P. van Schaik (Eds.): Orangutans: Geographic Variation in Behavioral Ecology and Conservation: Oxford University Press, pp. 119–134.

Ottoni, Eduardo B.; Resende, Briseida Dogo de; Izar, Patrícia (2005): Watching the best nutcrackers: what capuchin monkeys (Cebus apella) know about others’ tool-using skills. In Animal cognition 8 (4), pp. 215–219. DOI: 10.1007/s10071-004-0245-8.

Palagi, Elisabetta; Bergman, Thore J. (2021): Bridging Captive and Wild Studies: Behavioral Plasticity and Social Complexity in Theropithecus gelada. In Animals: an open access journal from MDPI 11 (10). DOI: 10.3390/ani11103003.

Perry, Susan; Jiminez, Juan Carlos Ordonez (2012): The effects of food size, rarity, and processing complexity on white-faced capuchins’ visual attention to foraging conspecifics. In Feeding ecology in apes and other primates 48, p. 203.

Pisor, Anne C.; Surbeck, Martin (2019): The evolution of intergroup tolerance in nonhuman primates and humans. In Evolutionary anthropology 28 (4), pp. 210–223. DOI: 10.1002/evan.21793.

Prasetyo, Didik; Ancrenaz, Marc; Morrogh-Bernard, Helen C.; Utami Atmoko, S. Suci; Wich, Serge A.; van Schaik, Carel P. (2009): Nest building in orangutans. In Serge A. Wich, S. Suci Utami Atmoko, Tatang Mitra Setia, Carel P. van Schaik (Eds.): Orangutans: Geographic Variation in Behavioral Ecology and Conservation: Oxford University Press, p. 0.

R Core Team (2023): R: A language and environment for statistical computing. Vienna, Austria: R Foundation for Statistical Computing. Available online at https://www.R-project.org/.

Rapaport, Lisa G.; Brown, Gillian R. (2008): Social influences on foraging behaviour in young nonhuman primates: Learning what, where, and how to eat. In Evolutionary Anthropology: Issues, News, and Reviews 17 (4), pp. 189–201. DOI: 10.1002/evan.20180.

Ross, Caroline; Jones, Kate E. (1999): Socioecology and the evolution of primate reproductive rates. In Comparative primate socioecology 22, pp. 73–110.

Schuppli, Caroline; Forss, Sofia; Meulman, Ellen; Atmoko, Suci Utami; van Noordwijk, Maria; van Schaik, Carel (2017): The effects of sociability on exploratory tendency and innovation repertoires in wild Sumatran and Bornean orangutans. In Scientific reports 7 (1), p. 15464. DOI: 10.1038/s41598-017-15640-x.

Schuppli, Caroline; Forss, Sofia I. F.; Meulman, Ellen J. M.; Zweifel, Nicole; Lee, Kevin C.; Rukmana, Evasari et al. (2016a): Development of foraging skills in two orangutan populations: needing to learn or needing to grow? In Frontiers in zoology 13, p. 43. DOI: 10.1186/s12983-016-0178-5.

Schuppli, Caroline; Isler, Karin; van Schaik, Carel P. (2012): How to explain the unusually late age at skill competence among humans. In Journal of Human Evolution 63 (6), pp. 843–850. DOI: 10.1016/j.jhevol.2012.08.009.

Schuppli, Caroline; Meulman, Ellen J.M.; Forss, Sofia I.F.; Aprilinayati, Fikty; van Noordwijk, Maria A.; van Schaik, Carel P. (2016b): Observational social learning and socially induced practice of routine skills in immature wild orang-utans. In Animal Behaviour 119, pp. 87–98. DOI: 10.1016/j.anbehav.2016.06.014.

Schuppli, Caroline; van Noordwijk, Maria; Atmoko, Suci Utami; van Schaik, Carel (2020): Early sociability fosters later exploratory tendency in wild immature orangutans. In Science advances 6 (2), eaaw2685. DOI: 10.1126/sciadv.aaw2685.

Schuppli, Caroline; van Schaik, Carel (2019a): Social learning among wild orang-utans: Is it affective? In Daniel Dukes, Fabrice Clément (Eds.): Foundations of Affective Social Learning: Conceptualizing the Social Transmission of Value. Cambridge: Cambridge University Press (Studies in Emotion and Social Interaction), pp. 25–40. Available online at https://www.cambridge.org/core/books/foundations-of-affective-social-learning/social-learning-among-wild-orangutans/1AA53F7E53E38B25431CE559C0195F94.

Schuppli, Caroline; van Schaik, Carel P. (2019b): Animal cultures: how we’ve only seen the tip of the iceberg. In Evolut. Hum. Sci. 1. DOI: 10.1017/ehs.2019.1.

Scott, Amy M.; Susanto, Tri Wayhu; Setia, Tatang Mitra; Knott, Cheryl D. (2023): Mother-offspring proximity maintenance as an infanticide avoidance strategy in bornean orangutans (Pongo pygmaeus wurmbii). In American Journal of Primatology 85 (6), e23482. DOI: 10.1002/ajp.23482.

Shumaker, Robert W.; Wich, Serge A.; Perkins, Lori (2008): Reproductive life history traits of female orangutans (Pongo spp.). In Interdisciplinary topics in gerontology 36, pp. 147–161. DOI: 10.1159/000137705.

Simpson, Gavin L. (2024): gratia: Graceful ggplot-Based Graphics and Other Functions for GAMs Fitted using mgcv. Available online at https://gavinsimpson.github.io/gratia/.

Stevens, Jeroen M. G.; Thierens, Marijke; Vervaecke, Hilde (2006): What are you looking at? The social contexts of peering behaviour in captive bonobos (Pan paniscus).

Stevens, Jeroen M. G.; Vervaecke, Hilde; Vries, Han de; van Elsacker, Linda (2005): Peering is not a formal indicator of subordination in bonobos (Pan paniscus). In American Journal of Primatology 65 (3), pp. 255–267. DOI: 10.1002/ajp.20113.

Tinbergen, N. (1963): On aims and methods of Ethology. In Zeitschrift für Tierpsychologie 20 (4), pp. 410–433. DOI: 10.1111/j.1439-0310.1963.tb01161.x.

van Noordwijk, M. A. (2012): From maternal investment to lifetime maternal care. In John C. Mitani, Josep Call, Peter M. Kappeler, Ryne A. Palombit, Joan B. Silk (Eds.): The evolution of primate societies. Chicago and London: The University of Chicago Press, pp. 321–342.

van Noordwijk, Maria A.; Atmoko, S. Suci Utami; Knott, Cheryl D.; Kuze, Noko; Morrogh-Bernard, Helen C.; Oram, Felicity et al. (2018): The slow ape: High infant survival and long interbirth intervals in wild orangutans. In Journal of Human Evolution 125, pp. 38–49. DOI: 10.1016/j.jhevol.2018.09.004.

van Noordwijk, Maria A.; LaBarge, Laura R.; Kunz, Julia A.; Marzec, Anna M.; Spillmann, Brigitte; Ackermann, Corinne et al. (2023): Reproductive success of Bornean orangutan males: scattered in time but clustered in space. In Behavioral ecology and sociobiology 77 (12), p. 134. DOI: 10.1007/s00265-023-03407-6.

van Noordwijk, Maria A.; Sauren, Simone E.B.; Abulani Ahbam, Nuzuar; Morrogh-Bernard, Helen C.; Utami Atmoko, S. Suci; van Schaik, Carel P. (2009): Development of independence: Sumatran and Bornean orangutans compared. In Serge A. Wich, S. Suci Utami Atmoko, Tatang Mitra Setia, Carel P. van Schaik (Eds.): Orangutans: Geographic Variation in Behavioral Ecology and Conservation: Oxford University Press, pp. 189–204.

van Schaik, C. P.; Burkart, J.; Damerius, L.; Forss, S. I. F.; Koops, K.; van Noordwijk, M. A.; Schuppli, C. (2016): The reluctant innovator: orangutans and the phylogeny of creativity. In Philosophical transactions of the Royal Society of London. Series B, Biological sciences 371 (1690). DOI: 10.1098/rstb.2015.0183.

van Schaik, Carel; Graber, Sereina; Schuppli, Caroline; Burkart, Judith (2017): The Ecology of Social Learning in Animals and its Link with Intelligence. In The Spanish journal of psychology 19, E99. DOI: 10.1017/sjp.2016.100.

van Schaik, Carel P. (1999): The socioecology of fission-fusion sociality in Orangutans. In Primates 40 (1), pp. 69–86. DOI: 10.1007/BF02557703.

van Schaik, Carel P.; Ancrenaz, Marc; Borgen, Gwendolyn; Galdikas, Birute; Knott, Cheryl D.; Singleton, Ian et al. (2003): Orangutan cultures and the evolution of material culture. In Science 299 (5603), pp. 102–105. DOI: 10.1126/science.1078004.

van Schaik, Carel P.; Burkart, Judith M. (2011): Social learning and evolution: the cultural intelligence hypothesis. In Philosophical transactions of the Royal Society of London. Series B, Biological sciences 366 (1567), pp. 1008–1016. DOI: 10.1098/rstb.2010.0304.

Vervaecke, H.; Vries, H. de; van Elsacker, L. (2000): Dominance and its Behavioral Measures in a Captive Group of Bonobos (Pan paniscus). In International Journal of Primatology 21 (1), pp. 47–68.

Watson, Stuart K.; Botting, Jennifer; Whiten, Andrew; van de Waal, Erica (2018): Culture and Selective Social Learning in Wild and Captive Primates. In Laura Desirèe Di Paolo, Fabio Di Vincenzo, Francesca de Petrillo (Eds.): Evolution of Primate Social Cognition, vol. 5. Cham: Springer International Publishing (Interdisciplinary Evolution Research), pp. 211–230.

Whiten, A.; Goodall, J.; McGrew, W. C.; Nishida, T.; Reynolds, V.; Sugiyama, Y. et al. (1999): Cultures in chimpanzees. In Nature 399, pp. 682–685. DOI: 10.1038/21415.

Whiten, Andrew; Horner, Victoria; Litchfield, Carla A.; Marshall-Pescini, Sarah (2004): How do apes ape? In Animal Learning & Behavior 32 (1), pp. 36–52. DOI: 10.3758/BF03196005.

Whiten, Andrew; van de Waal, Erica (2018): The pervasive role of social learning in primate lifetime development. In Behavioral ecology and sociobiology 72 (5), p. 80. DOI: 10.1007/s00265-018-2489-3.

Whiten, Andrew; van Schaik, Carel P. (2007): The evolution of animal ‘cultures’ and social intelligence. In Philosophical transactions of the Royal Society of London. Series B, Biological sciences 362 (1480), pp. 603–620. DOI: 10.1098/rstb.2006.1998.

Wich, Serge A.; Vries, Han de; Ancrenaz, Marc; Perkins, Lory; Shumaker, Robin W.; Suzuki, Akira; van Schaik, Carel P. (2009): Orangutan life history variation. In Serge A. Wich, S. Suci Utami Atmoko, Tatang Mitra Setia, Carel P. van Schaik (Eds.): Orangutans: Geographic Variation in Behavioral Ecology and Conservation: Oxford University Press, pp. 65–76.

Wood, S. N. (2017): Generalized Additive Models: An Introduction with R. 2nd ed.: Chapman and Hall/CRC.

Wood, S. N. (2018): Package ‘mgcv’ (R Package Version, 1.8-26). Available online at https://cran.r-project.org/web/packages/mgcv/.

