## SupplementaryMaterial for "The development of social attention in orangutans: comparing peering behaviour in wild and zoo-housed individuals"

**Supplementary material**

**Table S1. Overview table of all individuals included in this study.** The table shows the amount of data points per individual (N) in the full data sets, site of data collection, focal name, age classes observed, observation period, and observed age range.

| N | Site | Focal | Age class | Sex | Observation period | Age range (y) |
|---|---|---|---|---|---|---|
| 12 | Suaq | Alice | Adult | Female | 2010, 2017/18, 2020 | 50.5 – 59.5 |
| 5 | Suaq | Amor | Immature | Male | 2017/18 | 2.3 – 3.5 |
| 4 | Suaq | Balu | Adult | Male | 2017, 2019 | 24.7 – 26.8 |
| 7 | Suaq | Caesar | Adult | Male | 2018/19 | 21.6 – 23.1 |
| 12 | Suaq | Chindy | Immature | Female | 2008, 2010/11, 2017 | 5.8 – 14.6 |
| 61 | Suaq | Cinnamon | Immature | Female | 2013/14, 2016 – 2019 | 1.6 – 7.6 |
| 68 | Suaq | Cissy | Adult | Female | 2010/11, 2013/14, 2017 – 2019 | 45.8 – 54.3 |
| 4 | Suaq | Derek | Adult | Male | 2017 – 2019 | 23.6 – 25.3 |
| 2 | Suaq | Dian | Adult | Male | 2009, 2019 | 31.2 – 41.2 |
| 2 | Suaq | Diddy | Immature | Male | 2009, 2011 | 3.2 – 5.1 |
| 4 | Suaq | Dodi | Adult | Female | 2009, 2011, 2013 | 40.6 – 45.4 |
| 4 | Suaq | Dru | Adult | Male | 2018 | 24.3 – 24.4 |
| 68 | Suaq | Eden | Immature | Female | 2017 – 2020 | 2.2 – 5.2 |
| 92 | Suaq | Ellie | Adult | Female | 2009 – 2011, 2017 – 2020 | 17.9 – 20.9 |

| | | | | | | |
|---|---|---|---|---|---|---|
| 74 | Suaq | Frankie | Immature | Male | 2013/14, 2016 – 2019 | 0.9 – 7.3 |
| 30 | Suaq | Fredy | Immature | Male | 2008 – 2011, 2013/14 | 3.5 – 8.7 |
| 99 | Suaq | Friska | Adult | Female | 2008 – 2011, 2013/14, 2016 – 2019 | 65.9 – 76.9 |
| 4 | Suaq | Gura | Adult | Male | 2017/18 | 29.5 – 30.5 |
| 7 | Suaq | Islo | Adult | Male | 2008, 2018/19 | 31.2 – 42.3 |
| 2 | Suaq | Karma | Adult | Female | 2008, 2011 | 13.5 – 15.6 |
| 30 | Suaq | Lilly | Immature – Adult | Female | 2009, 2011, 2013, 2017 – 2019 | 7.9 – 18.0 |
| 115 | Suaq | Lisa | Adult | Female | 2008/9, 2011, 2013/14, 2016 – 2019 | 21.3 – 32.3 |
| 104 | Suaq | Lois | Immature | Male | 2011, 2013/14, 2016 – 2019 | 0.5 – 9.3 |
| 9 | Suaq | Luther | Immature | Male | 2017 – 2019 | 1.2 – 3.0 |
| 4 | Suaq | Marco | Adult | Male | 2018 | 20.8 |
| 1 | Suaq | Milo | Adult | Male | 2009 | 20.3 |
| 1 | Suaq | Otto | Adult | Male | 2017 | 40.0 |
| 7 | Suaq | Pepito | Immature | Male | 2017 – 2019 | 4.5 – 6.9 |
| 4 | Suaq | Pluto | Adult | Male | 2017 | 34.4 – 34.6 |

| | | | | | | |
|---|---|---|---|---|---|---|
| 12 | Suaq | Raffi | Adult | Female | 2014, 2017 | 41.2 – 44.6 |
| 11 | Suaq | Rendang | Immature | Male | 2014 | 0.7 – 0.9 |
| 1 | Suaq | Ronaldo | Immature | Male | 2011 | 5.0 |
| 4 | Suaq | Sazu | Immature | Male | 2014, 2017/18 | 7.6 – 11.9 |
| 8 | Suaq | Shera | Immature | Female | 2010/11 | 12.8 – 13.1 |
| 5 | Suaq | Simba | Immature | Male | 2014 | 0.9 |
| 9 | Suaq | Tina | Immature | Female | 2008, 2011 | 10.9 – 13.2 |
| 7 | Suaq | Tornado | Immature | Male | 2017 – 2019 | 2.8 – 4.7 |
| 10 | Suaq | Trident | Immature | Female | 2016 – 2019 | 10.4 – 13.2 |
| 1 | Suaq | Yoyo | Adult | Male | 2009 | 17.1 |
| 11 | Suaq | Yulia | Immature | Female | 2016, 2018 | 9.8 – 11.7 |
| 3 | Basel Zoo | Budi | Adult | Male | 2021 | 17.3 |
| 3 | Zurich Zoo | Cahaya | Adult | Female | 2021/22 | 19.1 – 20.2 |
| 8 | Dresden Zoo | Daisy | Adult | Female | 2021/22 | 30.2 – 31.6 |
| 12 | Dresden Zoo | Dalai | Immature | Male | 2021/2022 | 5.8 – 7.2 |
| 3 | Dresden Zoo | Djaka | Adult | Female | 2022 | 26.3 |
| 3 | Zurich Zoo | Djarius | Adult | Male | 2022 | 27.8 |

| | | | | | | |
|---|---|---|---|---|---|---|
| 3 | Dresden Zoo | Djudi | Adult | Female | 2022 | 49.3 |
| 3 | Zurich Zoo | Hadiah | Immature | Male | 2021 | 14 |
| 5 | Basel Zoo | Ketawa | Immature | Female | 2021 | 8.5 |
| 10 | Leipzig Zoo | Lursa | Immature | Female | 2022/23 | 0.6 – 1.5 |
| 3 | Basel Zoo | Maia | Adult | Female | 2021 | 13.8 |
| 6 | Zurich Zoo | Malou | Immature | Male | 2021/22 | 9.7 – 10.4 |
| 6 | Zurich Zoo | Mimpi | Immature | Female | 2021/22 | 9.3 – 10.0 |
| 3 | Basel Zoo | Ombak | Immature | Male | 2022 | 5.3 |
| 5 | Leipzig Zoo | Padana | Adult | Female | 2021 | 23.5 – 23.7 |
| 6 | Basel Zoo | Padma | Immature | Female | 2021/23 | 3.1 – 4.6 |
| 12 | Zurich Zoo | Pandai | Immature | Female | 2021 - 2023 | 6.0 – 7.9 |
| 3 | Leipzig Zoo | Raja | Adult | Female | 2022 | 19.0 |
| 6 | Basel Zoo | Revital | Adult | Female | 2021 | 21.2 – 21.3 |
| 9 | Zurich Zoo | Riang | Immature | Female | 2021/22 | 4.0 – 5.5 |
| 15 | Leipzig Zoo | Sari | Immature | Female | 2021 – 2023 | 3.8 – 5.6 |
| 2 | Zurich Zoo | Timor | Adult | Female | 2022 | 46.8 |
| 3 | Dresden Zoo | Toni | Adult | Male | 2021 | 29.6 |
| 11 | Zurich Zoo | Utu | Immature | Female | 2021/22 | 1.3 – 2.8 |

| | | | | | | |
|---|---|---|---|---|---|---|
| 3 | Zurich Zoo | Xira | Adult | Female | 2023 | 25.6 |

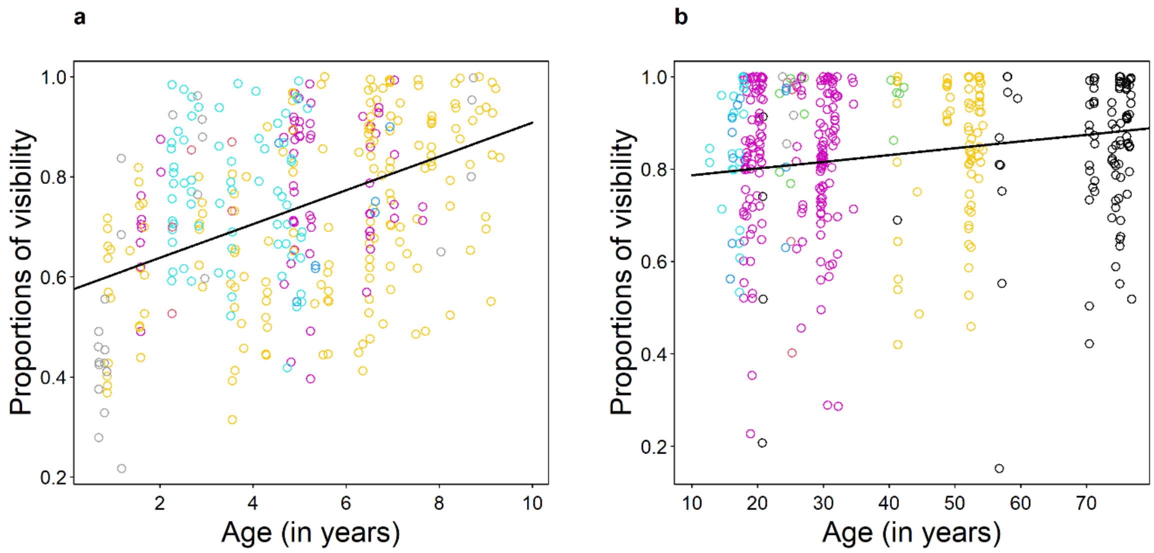


**Figure S1. Proportion of focal visibility over age in the wild population.** Proportion of visible observation time of each focal follow day in Suaq, Indonesia over age for (a) 0 to 10 years, and (b) 10+ years. Each point represents one focal follow day. Colours refer to different individuals. The lines represent mean model predictions (plotted using the *ggpredict* function of the ggeffects package (Lüdecke 2018)).

**Table S2. Model summary for model M1a+b.** Linear Mixed-Effects Model for focal visibility in the wild data set. Analysis conducted using the *lmer* function of the lme4 package (Bates et al. 2015). The table shows estimates, standard errors, P values, sample size (N) and confidence intervals. P values with significance at the 5 % level are indicated in bold font (assessed with the lmerTest package, Kuznetsova et al. 2017). CI = confidence interval.

| Model | Dependent variable | Effect | Effect type | Estimate | Std. Error | P value | 95 % CI |
|---|---|---|---|---|---|---|---|
| M1a<br><br>n = 344 | Proportion of visibility (0 – 10 years) | (Intercept)<br>Age<br>Individual ID | Intercept<br>Fixed effect<br>Random effect | 0.571<br>0.038<br>- | 0.030<br>0.004<br>- | **<0.001**<br>**<0.001**<br>- | 0.512 – 0.629<br>0.025 – 0.042<br>- |
| M1b<br><br>n = 423 | Proportion of visibility (10+ years) | (Intercept)<br>Age<br>Individual ID | Intercept<br>Fixed effect<br>Random effect | 0.772<br>0.001<br>- | 0.038<br>0.001<br>- | **<0.001**<br>0.186<br>- | 0.697 – 0.846<br>-0.001 – 0.003 |

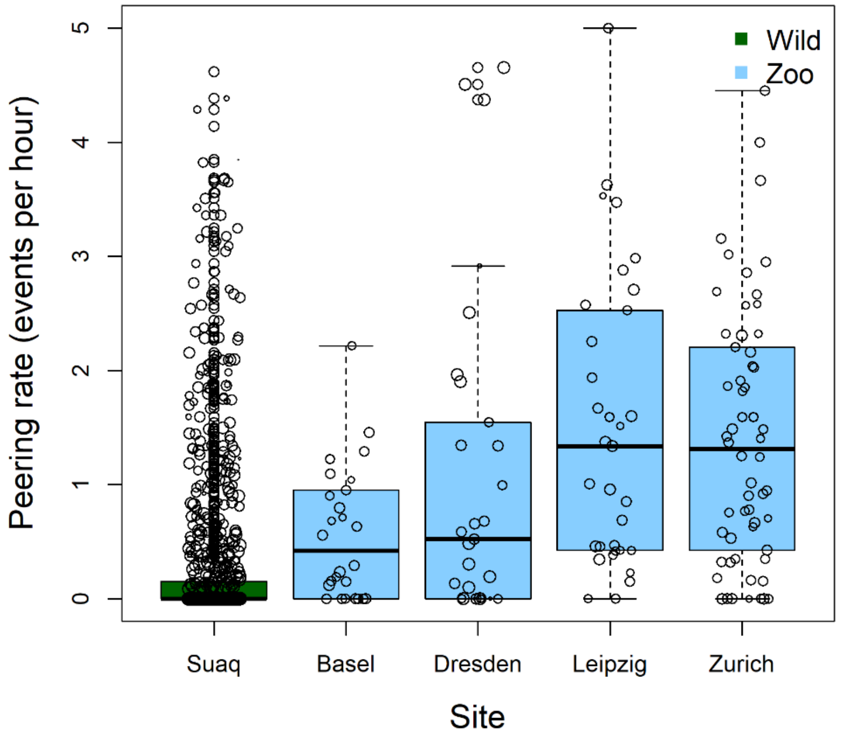


**Figure S2. Peering frequency per site.** Mean hourly peering rates across the five sites of data collection. The bold horizontal line indicates the median, the box the lower and upper quartile, the whiskers the lowest and greatest value, respectively, and circles are outliers. Each point represents one focal follow day.

**Table S3. Development of peering frequency over age: summary for model 2c.** GAMM with site (Suaq, Basel, Dresden, Leipzig, Zurich), age (z-transformed), as well as an interaction between age and site as fixed effects, and individual and observer ID as random effects. The model included visible observation duration as an offset term. Listed are estimates, standard errors, p values, sample size (n), adjusted $r^2$, and deviance explained (DE). P values with significance at the 5 % level are indicated in bold font.

| Model | Response variable | Predictors | Type | Estimate | Standard Error | p-value |
|---|---|---|---|---|---|---|
| 2c | Peering | <u>Parametric terms:</u> | | | | |
| | counts | (Intercept) | Intercept | -3.308 | 0.239 | **< 0.001** |
| n = 1061 | total (all | Site (Zoo Basel) | Fixed effect | 0.032 | 0.677 | 0.963 |
| $r^2_{adj.}$ = 0.63 | ages) | Site (Zoo Dresden) | Fixed effect | 2.634 | 0.374 | **< 0.001** |
| DE = | | Site (Zoo Leipzig) | Fixed effect | 3.463 | 5.243 | 0.509 |
| 73.7% | | Site (Zoo Zürich) | Fixed effect | 2.173 | 0.404 | **< 0.001** |

| | | | | | | |
|---|---|---|---|---|---|---|
| | | <u>Smooth terms:</u> | | | | |
| | | Z-Age | Fixed effect | | | **< 0.001** |
| | | Z-Age:Suaq | Interaction | | | **< 0.001** |
| | | Z-Age:Zoo Basel | Interaction | | | 0.998 |
| | | Z-Age:Zoo Dresden | Interaction | | | **0.014** |
| | | Z-Age:Zoo Leipzig | Interaction | | | **0.020** |
| | | Z-Age:Zoo Zurich | Interaction | | | 0.065 |
| | | Individual ID | Random effect | | | **0.002** |
| | | Observer ID | Random effect | | | **< 0.001** |

**Table S4. Comparison between peering frequency per sites**. Post-hoc pairwise comparison of estimated marginal means of the data collection sites (Model S2), using the *emmeans* function from the emmeans package (Lenth 2023). The table shows contrasts, estimates, standard errors, t-ratio, and p values for each comparison. P values with significance at the 5 % level are indicated in bold font.

| Contrast | Estimate | Standard Error | t-ratio | p-value |
|---|---|---|---|---|
| Suaq – Zoo Basel | -1.694 | 0.719 | -2.356 | 0.129 |
| Suaq – Zoo Dresden | -4.223 | 0.455 | -9.284 | **< 0.001** |
| Suaq – Zoo Leipzig | -4.085 | 0.554 | -7.377 | **< 0.001** |
| Suaq – Zoo Zürich | -3.835 | 0.473 | -8.114 | **< 0.001** |
| Zoo Basel – Zoo Dresden | -2.529 | 0.706 | -3.582 | 0.003 |
| Zoo Basel – Zoo Leipzig | -2.391 | 0.785 | -3.046 | 0.020 |
| Zoo Basel – Zoo Zürich | -2.142 | 0.719 | -2.978 | 0.025 |
| Zoo Dresden – Zoo Leipzig | 0.138 | 0.553 | 0.250 | 1.000 |
| Zoo Dresden – Zoo Zürich | 0.388 | 0.445 | 0.871 | 0.908 |
| Leipzig Zoo – Zoo Zürich | 0.250 | 0.558 | 0.448 | 0.992 |

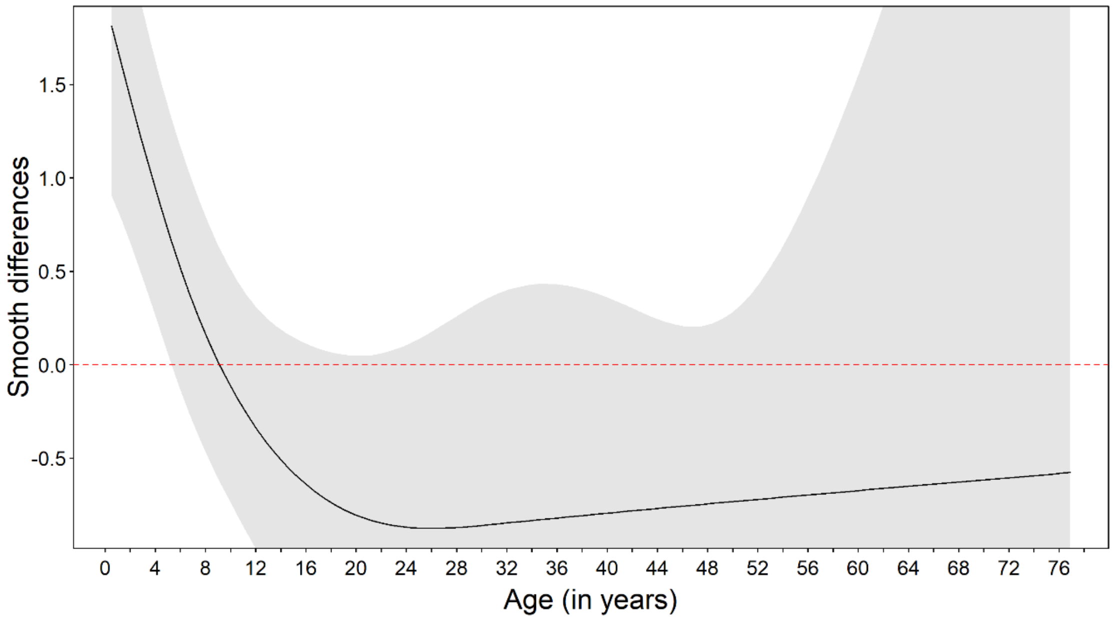


**Figure S3. Differences in age trends in peering frequency.** Differences between the smoothing splines of the two settings in Model 2a for peering in the wild and in the zoos over age. Grey areas are 95 % confidence intervals. Areas of statistical significance at the 5 % level are assesses by whether or where y = 0 (red dotted line) crosses the confidence interval.

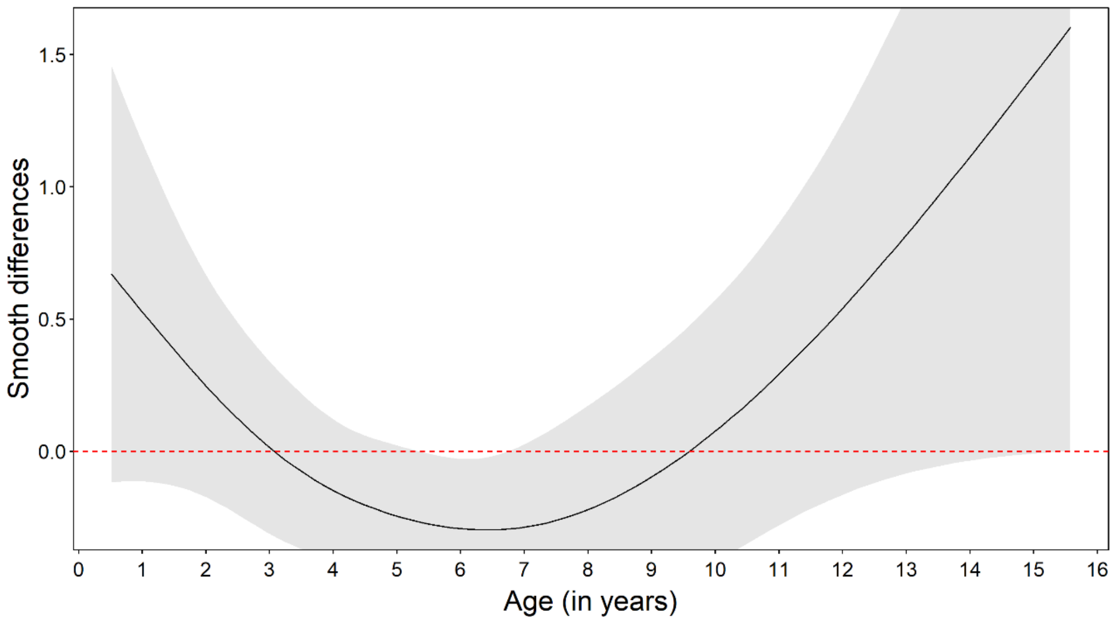


**Figure S4. Differences in age trends in peering frequency controlled for the availability of association partners.** Differences between the smoothing splines of the two settings in Model 2b for peering in peering range in the wild and the zoos over age. Grey areas are 95 % confidence intervals. Areas of statistical significance at the 5 % level are assesses by whether or where y = 0 (red dotted line) crosses the confidence interval.

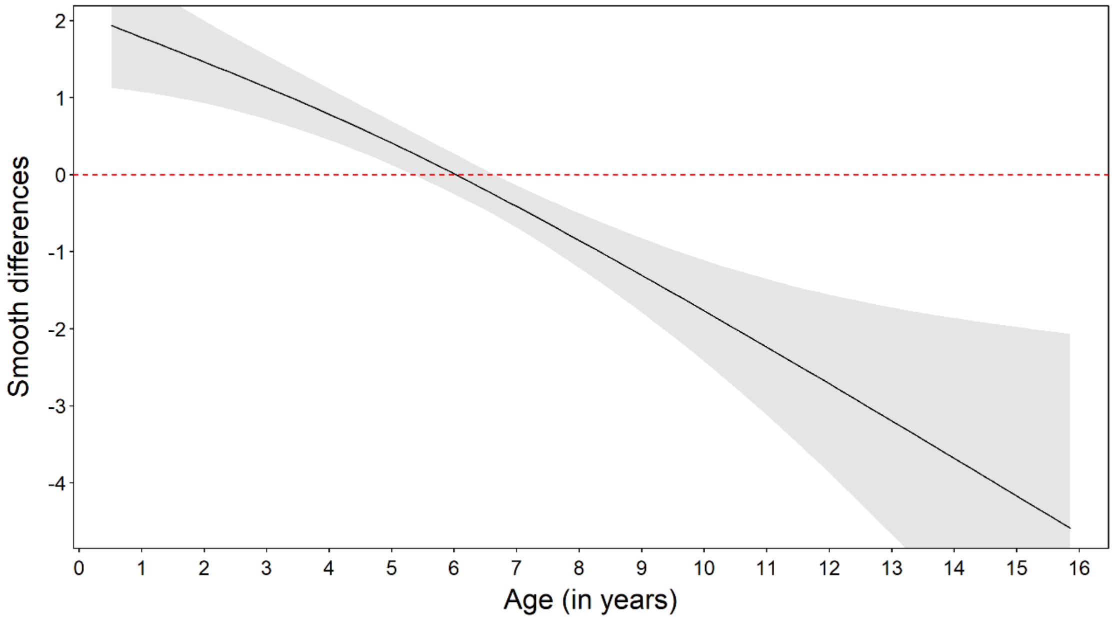


**Figure S5. Differences in age trends in peering frequency at the mother.** Differences between the smoothing splines of the two settings in Model 3a for peering directed at the mother in the wild and the zoos over age. Grey areas are 95 % confidence intervals. Areas of statistical significance at the 5 % level are assesses by whether or where y = 0 (red dotted line) crosses the confidence interval.

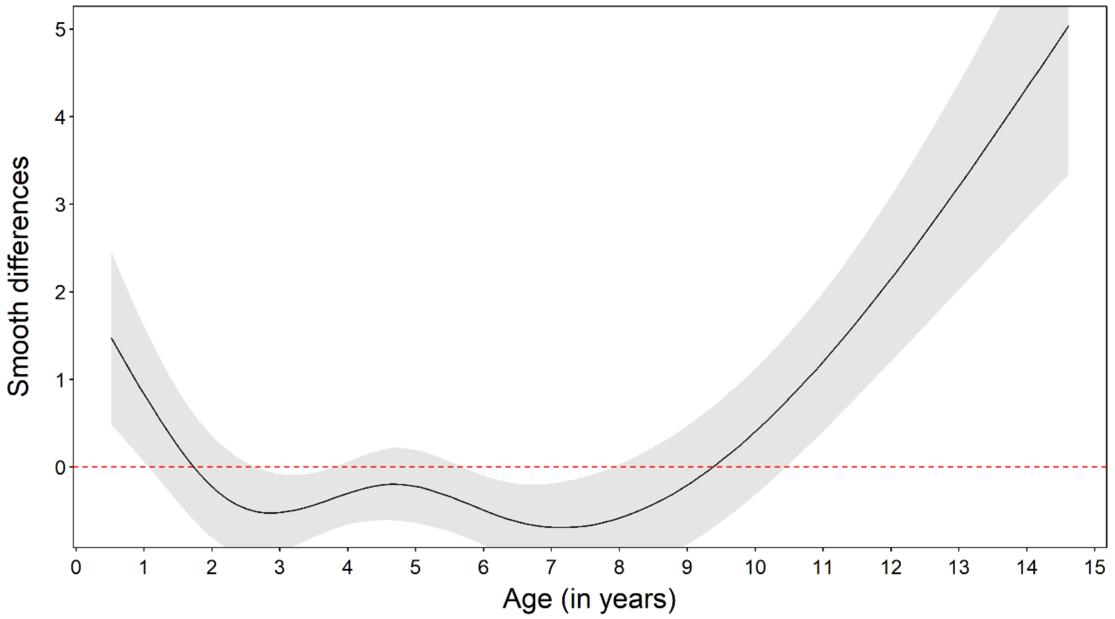


**Figure S6. Differences in age trends for peering frequencies at non-mother targets controlled for opportunities to peer.** Differences between the smoothing splines of the two settings in Model 3c for peering proportions directed at non-mother individuals in the wild and the zoos over age. Grey areas are 95 % confidence intervals. Areas of statistical significance at the 5 % level are assesses by whether or where y = 0 (red dotted line) crosses the confidence interval.

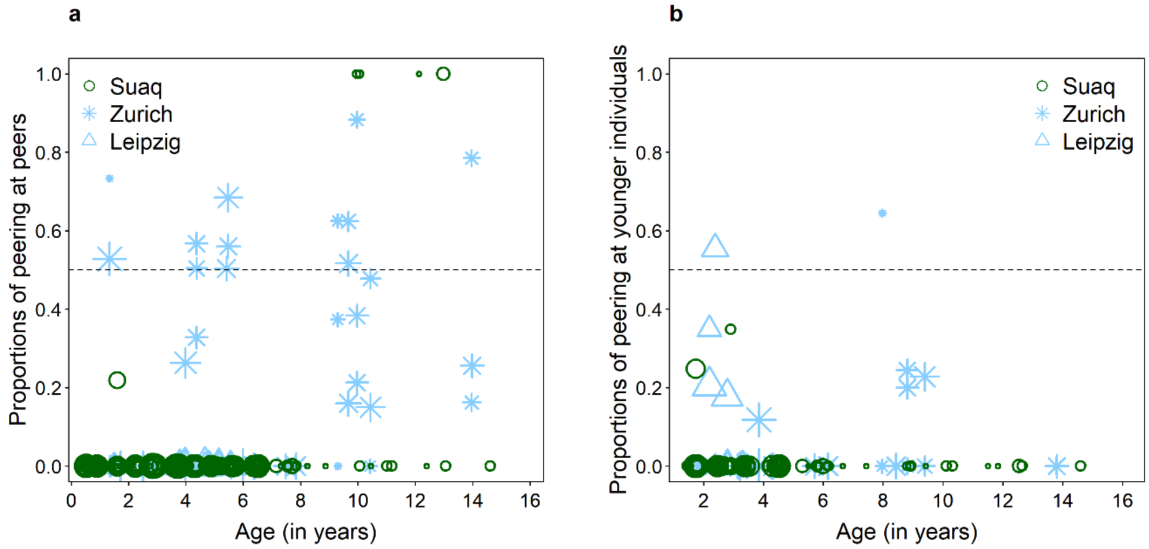


**Figure S7. Development of peering frequencies at peering targets of different relative ages controlled for opportunities to peer.** Proportions of peering by wild and zoo-housed immature orangutans directed at (a) peers, and (b) younger individuals controlled for the time the peering individual spent in close proximity to the peering targets. Each point represents one focal follow day. Symbol sizes correspond to the log values of the total number of observed peering events.

**Table S5. Peering target selection: summary of models S3e and 3f.** GAMMs with setting (Wild / Zoo), age (z-transformed), as well as an interaction between age and setting as fixed effects, and individual and observer ID as random effects. Listed are estimates, standard errors, p values, sample size (n), adjusted $r^2$, and deviance explained (DE). P values with significance at the 5 % level are indicated in bold font.

| Model | Response variable | Predictors | Type | Estimate | Standard Error | p-value |
|---|---|---|---|---|---|---|
| 3e | Proportions | Parametric terms: | | | | |
| | of peering | (Intercept) | Intercept | -2.663 | 0.285 | **< 0.001** |
| n = 308 | at peers | Setting (zoo) | Fixed effect | 0.915 | 0.501 | 0.068 |
| $r^2_{adj.}$ = | (immatures) | Smooth terms: | | | | |
| 0.59 | | Z-Age | Fixed effect | | | **< 0.001** |
| DE = 71.4 | | Z-Age:Wild | Interaction | | | 1.000 |
| % | | Z-Age:Zoo | Interaction | | | 0.112 |
| | | Individual ID | Random effect | | | **< 0.001** |
| | | Observer ID | Random effect | | | **< 0.001** |
| 3f | Proportions | Parametric terms: | | | | |
| | of peering | (Intercept) | Intercept | -2.614 | 0.122 | **< 0.001** |
| n = 167 | at younger | Z-Age | Fixed effect | 0.027 | 0.046 | 0.557 |
| $r^2_{adj.}$ = | individuals | Setting (zoo) | Fixed effect | 0.361 | 0.156 | **0.021** |
| 0.12 | (immatures) | Z-Age:Zoo | Interaction | -0.014 | 0.091 | 0.880 |
| DE = 79 % | | Smooth terms: | | | | |
| | | Individual ID | Random effect | | | 0.455 |
| | | Observer ID | Random effect | | | **< 0.001** |

**Table S6 Context selection: summaries for models 4a-j.** GAMMs with setting (Wild/Zoo), age (z-transformed), as well as an interaction between age and setting as fixed effects, and individual and observer ID as random effects. Models 4f-j included the time association partners spent engaging in the behaviour as the offset term. Listed are estimates, standard errors, p values, sample size (n), adjusted $r^2$, and deviance explained (DE). P values with significance at the 5 % level and corrected for multiple testing due to the dependence in the response variable are indicated in bold font.

| Model | Response variable | Predictors | Type | Estimate | Standard Error | p-value |
|---|---|---|---|---|---|---|
| 4a | Proportion | Parametric terms: | | | | |
| | of peering | (Intercept) | Intercept | 1.881 | 0.427 | **< 0.001** |
| n = 246 | at feeding | Setting (zoo) | Fixed effect | -1.931 | 0.631 | **0.002** |
| $r^2_{adj.}$ = | behavior | Smooth terms: | | | | |
| 0.29 | (dependent | Z-Age | Fixed effect | | | **< 0.001** |
| DE = | immatures) | Z-Age:Wild | Interaction | | | **0.004** |
| 51.8 % | | Z-Age:Zoo | Interaction | | | 0.940 |
| | | Individual ID | Random effect | | | **< 0.001** |
| | | Observer ID | Random effect | | | **< 0.001** |
| 4b | Proportion | Parametric terms: | | | | |
| | of peering | (Intercept) | Intercept | -1.112 | 0.175 | **< 0.001** |
| n = 246 | at nesting | Setting (zoo) | Fixed effect | -0.682 | 0.252 | **0.007** |
| $r^2_{adj.}$ = | behavior | Smooth terms: | | | | |
| 0.10 | (dependent | Z-Age | Fixed effect | | | **< 0.001** |
| DE = | immatures) | Z-Age:Wild | Interaction | | | 0.149 |
| 55.5 % | | Z-Age:Zoo | Interaction | | | 0.900 |
| | | Individual ID | Random effect | | | **< 0.001** |
| | | Observer ID | Random effect | | | **0.002** |
| 4c | Proportion | Parametric terms: | | | | |
| | of peering | (Intercept) | Intercept | -3.927 | 0.184 | **< 0.001** |
| n = 246 | at social | Z-Age | Fixed effect | -0.020 | 0.044 | 0.648 |
| $r^2_{adj.}$ = | behavior | Setting (zoo) | Fixed effect | 0.839 | 0.252 | **< 0.001** |
| 0.02 | (dependent | Smooth terms: | | | | |
| DE = | immatures) | Z-Age:Wild | Interaction | | | 0.205 |
| 29.3 % | | Z-Age:Zoo | Interaction | | | **< 0.001** |
| | | Individual ID | Random effect | | | **< 0.001** |
| | | Observer ID | Random effect | | | **0.011** |
| 4d | Proportion | Parametric terms: | | | | |
| | of peering | (Intercept) | Intercept | -2.758 | 0.215 | **< 0.001** |
| n = 246 | at | Setting (zoo) | Fixed effect | 1.750 | 0.308 | **< 0.001** |
| $r^2_{adj.}$ = | exploratory | Smooth terms: | | | | |
| 0.49 | behavior | Z-Age | Fixed effect | | | **< 0.001** |

| | | | | | | |
|---|---|---|---|---|---|---|
| DE = 59 | (dependent | Z-Age:Wild | Interaction | | | **0.009** |
| % | immatures) | Z-Age:Zoo | Interaction | | | **0.025** |
| | | Individual ID | Random effect | | | **< 0.001** |
| | | Observer ID | Random effect | | | **< 0.001** |
| 4e | Proportion | Parametric terms: | | | | |
| | of peering | (Intercept) | Intercept | -4.079 | 0.191 | **< 0.001** |
| n = 246 | at other | Setting (zoo) | Fixed effect | 1.200 | 0.175 | **< 0.001** |
| $r^2_{adj.}$ = | (non- | Smooth terms: | | | | |
| 0.13 | social) | Z-Age | Fixed effect | | | **0.045** |
| DE = | behavior | Z-Age:Wild | Interaction | | | 0.166 |
| 49.9 % | (dependent | Z-Age:Zoo | Interaction | | | **< 0.001** |
| | immatures) | Individual ID | Random effect | | | **0.033** |
| | | Observer ID | Random effect | | | **< 0.001** |
| 4f | Peering | Parametric terms: | | | | |
| | counts at | (Intercept) | Intercept | -1.257 | 0.270 | **< 0.001** |
| n = 378 | feeding | Setting (zoo) | Fixed effect | 1.560 | 0.285 | **< 0.001** |
| $r^2_{adj.}$ = | behavior | Smooth terms: | | | | |
| 0.53 | (dependent | Z-Age | Fixed effect | | | **0.001** |
| DE = | immatures) | Z-Age:Wild | Interaction | | | **0.001** |
| 64.1 % | | Z-Age:Zoo | Interaction | | | 0.984 |
| | | Individual ID | Random effect | | | **< 0.001** |
| | | Observer ID | Random effect | | | **< 0.001** |
| 4g | Peering | Parametric terms: | | | | |
| | counts at | (Intercept) | Intercept | 0.102 | 0.324 | 0.734 |
| n = 323 | nesting | Z-Age | Fixed effect | -0.289 | 0.112 | **0.010** |
| $r^2_{adj.}$ = | Behavior | Setting (zoo) | Fixed effect | -0.103 | 0.492 | 0.834 |
| 0.31 | (dependent | Z-Age:Setting (zoo) | Interaction | 0.513 | 0.388 | 0.186 |
| DE = | immatures) | Smooth terms: | | | | |
| 40.6 % | | Individual ID | Random effect | | | 0.295 |
| | | Observer ID | Random effect | | | **< 0.001** |
| 4h | Peering | Parametric terms: | | | | |
| | counts at | (Intercept) | Intercept | -1.753 | 0.486 | **< 0.001** |
| n = 240 | social | Z-Age | Fixed effect | -0.403 | 0.395 | 0.307 |
| | | Setting (zoo) | Fixed effect | 0.603 | 0.543 | 0.267 |

| | | | | | | |
|---|---|---|---|---|---|---|
| $r^2_{adj.}$ = | Behavior | Z-Age:Setting(zoo) | Interaction | 0.417 | 0.900 | 0.368 |
| 0.12 | (dependent | Smooth terms: | | | | |
| DE = | immatures) | Individual ID | Random effect | | | 0.342 |
| 22.6 % | | Observer ID | Random effect | | | **0.012** |
| 4i | Peering | Parametric terms: | | | | |
| | counts at | (Intercept) | Intercept | -0.063 | 0.721 | 0.930 |
| n = 119 | exploratory | Z-Age | Fixed effect | 0.456 | 0.790 | 0.564 |
| $r^2_{adj.}$ = | Behavior | Setting (zoo) | Fixed effect | 0.163 | 0.745 | 0.827 |
| 0.44 | (dependent | Z-Age:Setting(zoo) | Interaction | -0.321 | 0.809 | 0.691 |
| DE = | immatures) | Smooth terms: | | | | |
| 16.2 % | | Individual ID | Random effect | | | **0.016** |
| | | Observer ID | Random effect | | | 0.285 |
| 4j | Peering | Parametric terms: | | | | |
| | counts at | (Intercept) | Intercept | -5.454 | 0.717 | **< 0.001** |
| n = 378 | other (non- | Setting (zoo) | Fixed effect | 2.643 | 0.962 | **0.006** |
| $r^2_{adj.}$ = | social) | Smooth terms: | | | | |
| 0.47 | Behavior | Z-Age | Fixed effect | | | **< 0.001** |
| DE = | (dependent | Z-Age:Wild | Interaction | | | **0.042** |
| 69.7 % | immatures) | Z-Age:Zoo | Interaction | | | **0.008** |
| | | Individual ID | Random effect | | | **0.002** |
| | | Observer ID | Random effect | | | **0.008** |

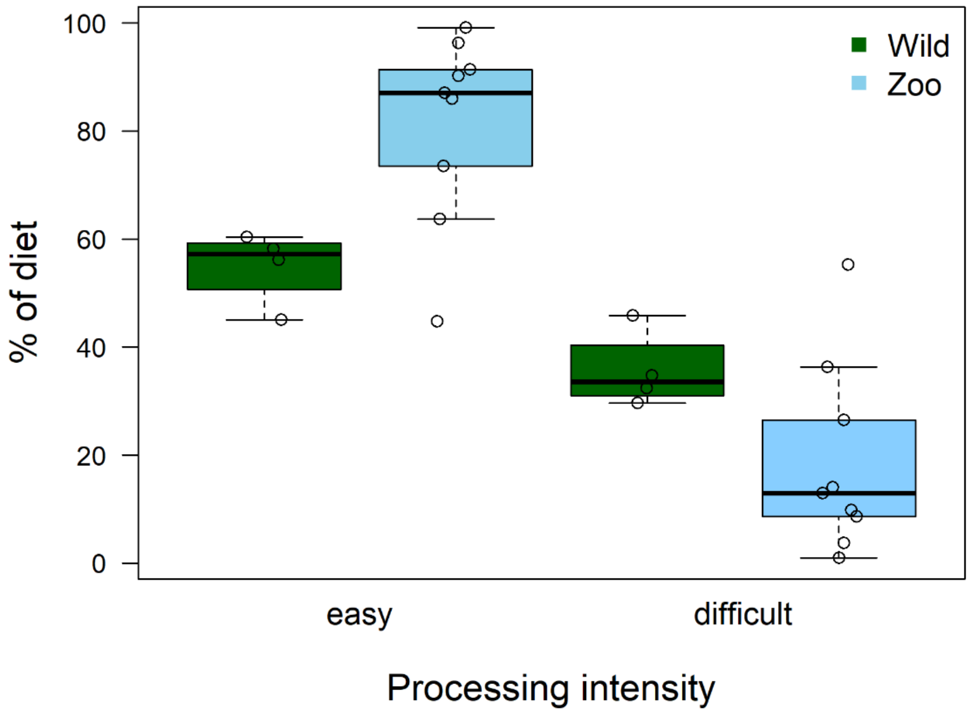


**Figure S8. Comparison of processing intensity of the two settings.** Percentage of easy and difficult food items in the diet of adult wild and zoo-housed orangutans. Based on data of 4 adult females in at Suaq, and of 9 adult individuals (2 males and 7 females) in the zoos. Percentages were calculated based on time spent feeding on the different item types.

### References


Bates, D.; Mächler, M.; Bolker, B.; Walker, S. (2015): Fitting Linear Mixed-Effects Models Using lme4. In *Journal of Statistical Software* 67 (1), pp. 1–48. DOI: 10.18637/jss.v067.i01.

Kuznetsova, A.; Brockhoff, P. B.; Christensen, R. H. B. (2017): lmerTest Package: Tests in Linear Mixed Effects Models. In *Journal of Statistical Software* 82 (13), pp. 1–26. DOI: 10.18637/jss.v082.i13.

Lenth, Russel V. (2023): emmeans: Estimated Marginal Means, aka Least-Squares Means. Available online at https://CRAN.R-project.org/package=emmeans.

Lüdecke, D. (2018): ggeffects: Tidy Data Frames of Marginal Effects from Regression Models. In *Journal of Open Source Software* 3 (26), p. 772. DOI: 10.21105/joss.00772.
